# A unified theory for predicting pathogen mutual invasibility and co-circulation

**DOI:** 10.1101/2024.04.22.590623

**Authors:** Sang Woo Park, Sarah Cobey, C. Jessica E. Metcalf, Jonathan M. Levine, Bryan T. Grenfell

## Abstract

A key aim in the dynamics and control of infectious diseases is predicting competitive outcomes of pathogen interactions. Observed pathogen community structure indicates both considerable coexistence of related variants and spectacular instances of replacement, notably in seasonal influenza and SARS-CoV-2. However, an overall comparative quantitative framework for invasion and coexistence remains elusive. Inspired by modern ecological coexistence theory, we address this gap by developing pathogen invasion theory (PIT) and test the resulting framework against empirical systems. PIT predicts near-universal mutual invasibility across major pathogen systems, including seasonal influenza strains and SARS-CoV-2 variants. Predicting co-circulation from mutual invasibility further depends on the extent of overcompensatory susceptible depletion dynamics. Our analyses highlight the central role of immuno-epidemiological factors in determining pathogen coexistence and community structure.

## Introduction

Understanding the determinants of coexistence or exclusion of pathogen variants is a fundamental question in the dynamics and control of infectious diseases (*1–6*). More than 1300 species of human pathogens have been identified so far (*7*), many responsible for substantial public health burdens. However, it is unclear why so many (or so few) can stably coexist in human populations (*5*). Despite many studies on particular host-pathogen systems (*8–13*), there is no overall comparative conceptual framework for explaining invasion dynamics and subsequent competitive outcomes across pathogen systems. The continued emergence of novel pathogens and ongoing vaccine development against antigenically variable pathogens stresses the importance of addressing this gap (*14*).

A major challenge in developing a unifying framework for pathogen community structure is to explain the heterogeneity of observed competition. Specifically, what mechanisms promote stable coexistence among some pathogen variants (Fig. 1A–C) but not among others (Fig. 1D–E)? Even among stably coexisting pathogen variants, there is a wide heterogeneity in the observed dynamics. For example, RSV A and B exhibit cyclic patterns of dominance (Fig. 1A), whereas the relative abundances of Coxsackievirus A16 and Enterovirus A71 remain roughly stable over time (Fig. 1B). Moreover, the coexistence patterns of four dengue serotypes seem to share both of these features (Fig. 1C). On the other hand, the evolutionary dynamics of SARS-CoV-2 variants (Fig. 1D) and seasonal influenza strains of the same subtype (Fig. 1E) exemplify permanent strain replacements, where the emergence of antigenically distinct variants causes extinction of previously dominant variants (*8,15*). Explaining and predicting competitive outcomes across pathogen systems remain a key aim.

**Figure 1.**
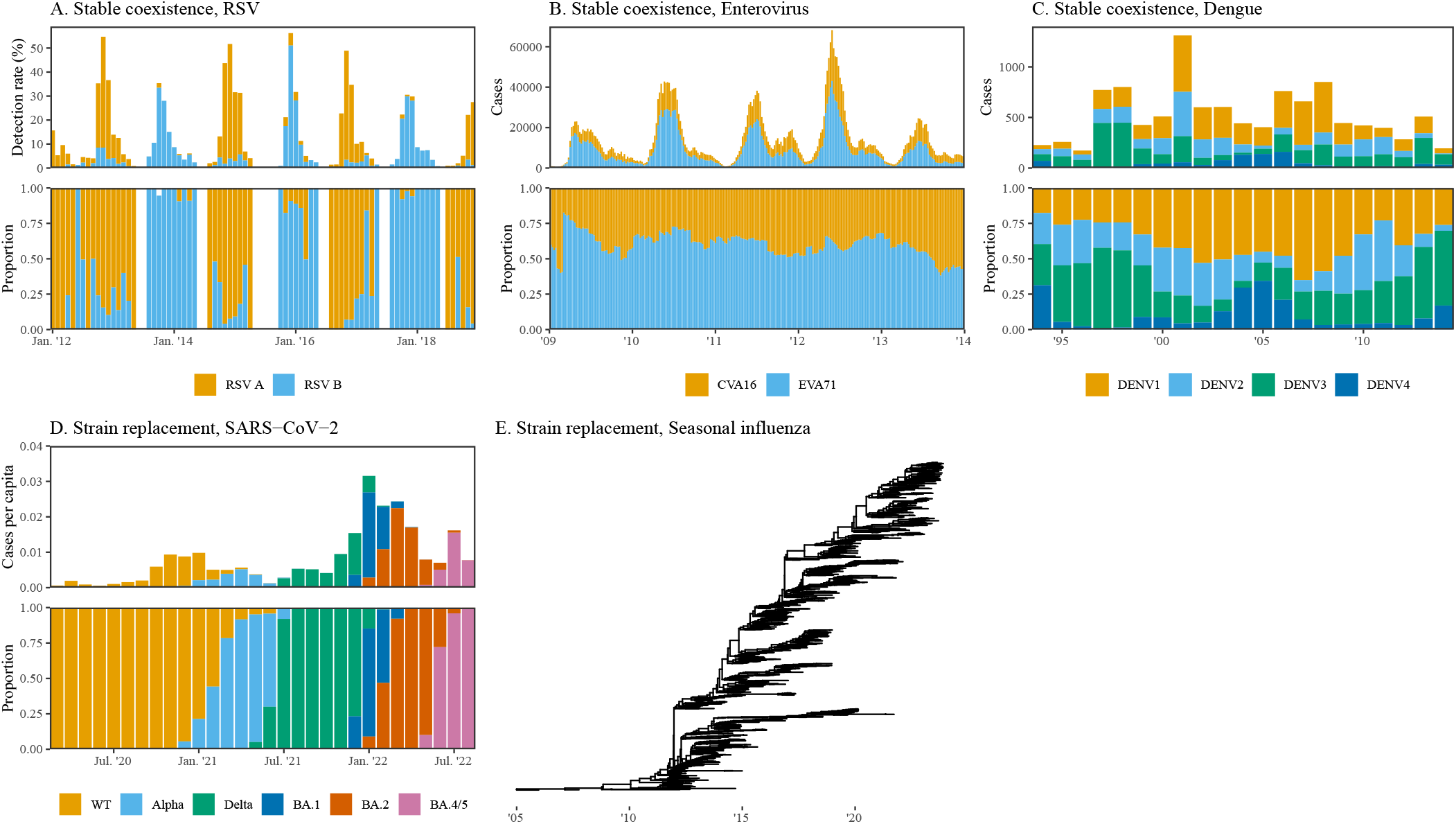
Observed coexistence patterns of human pathogens. (A) Monthly detection rates of RSV A and B (top) and their relative proportions (bottom) from Cheonan, Korea (*16*). (B) Weekly cases of hand, foot, and mouth disease caused by CV-A16 and EV-A71 (top) and their relative proportions (bottom) from all 31 provinces of Mainland China (*17*). (C) Annual cases of four dengue serotypes (top) and their relative proportions (bottom) from Bangkok, Thailand (*18*). (D) Monthly per capita cases of SARS-CoV-2 variants (top) and their relative proportions (bottom) from 10 representative countries (*15*). (E) Phylogenetic tree of 1578 human influenza A H3N2 subtype genome sampled between 2005 and 2023 from https://nextstrain.org/. All data are taken from publicly available sources from above references.

### Pathogen Invasion Theory

Many models have been proposed to explain the observed patterns of competition for specific pathogen systems (*10, 11, 19, 20*) and to further identify mechanisms that dictate persistence and coexistence following invasion of new variants (*21, 22*). However, these models often rely on detailed assumptions, often specific to the pathogen, making it difficult to draw general conclusions about pathogen competition across different systems. So far, there is no single, overarching framework that allows quantitative comparisons of coexistence potential across different pathogen systems.

To unify heterogeneous patterns of pathogen competition, we develop a pathogen invasion theory (PIT), inspired by Chesson’s modern coexistence theory (MCT) from community ecology (Section S1; (*23*)). Specifically, PIT allows us to boil down any system of two competing pathogen variants *i* and *j* into two fundamental quantities that determine their invasibility (Section S2–S3). First, the **immunological niche difference** 1−ρ describes the ability of a variant to escape the host’s immune response to another variant. This is measured as a ratio of equilibrium values (or long-term averages) of intraspecific susceptibility (*S*_*i*|*i*_ and *S*_*j*|*j*_) versus interspecific susceptibility (*S*_*j*|*i*_ and *S*_*i*|*j*_) at the population level:

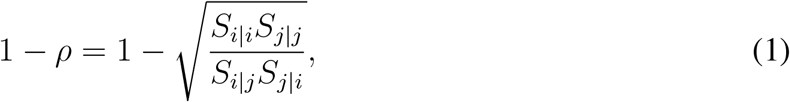

where *S*_*j*|*i*_ represent the effective proportion of individuals who are susceptible to variant *j* when variant *i* is at equilibrium; note that equilibrium conditions arise when ℛ_0,*i*_*S*_*i*|*i*_ = 1. Second, the **fitness difference**, *f*_*i*_*/f*_*j*_, measures the ratio of each variant’s innate ability to spread, discounted by its susceptibility to the host immune response to the other variant:

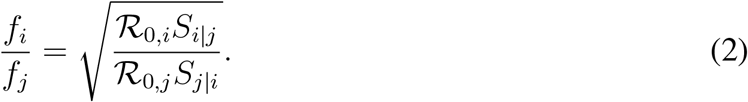

Then, like MCT (*23*), the immunological niche difference bounds the fitness differences compatible with mutual invasion. In short, the greater niche difference 1 − *ρ* (likewise, the smaller niche overlap *ρ*), the more easily competitively imbalanced strains can mutually invade, satisfying the following inequality:

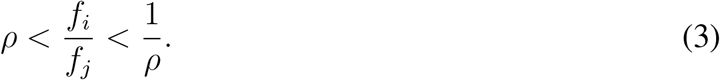

Importantly, all of the quantities necessary to inform the immunological niche difference and the fitness difference *(ℛ*_0,*i*_, *S*_*i*|*j*_) can be derived from mechanistic epidemiological models tailored to individual diseases, allowing for quantitative comparisons across disease systems.

Mutual invasion is often associated with long-term, stable coexistence in the classical MCT rooted in competitive interactions (*23*). However, nonlinear host-pathogen interactions, especially immunological feedback, add challenges to translating mutual invasibility to predictions about prolonged pathogen co-circulation. Here, we begin by comparing invasibility across pathogen systems and then turn to predicting pathogen co-circulation from invasibility.

#### Comparative pathogen invasion dynamics

We quantify mutual invasibility of various pathogen variant pairs by applying PIT (Section S4) to a diverse set of empirically motivated models of multistrain dynamics in various pathogens (*1, 14, 15, 19, 24–29*). The results are plotted in Fig. 2, where the region of mutual invasibility is bounded by Eq. (3). The boundaries of mutual invasibility, predicted by PIT, are equivalent to those predicted for ecological communities by MCT (*23*).

**Figure 2.**
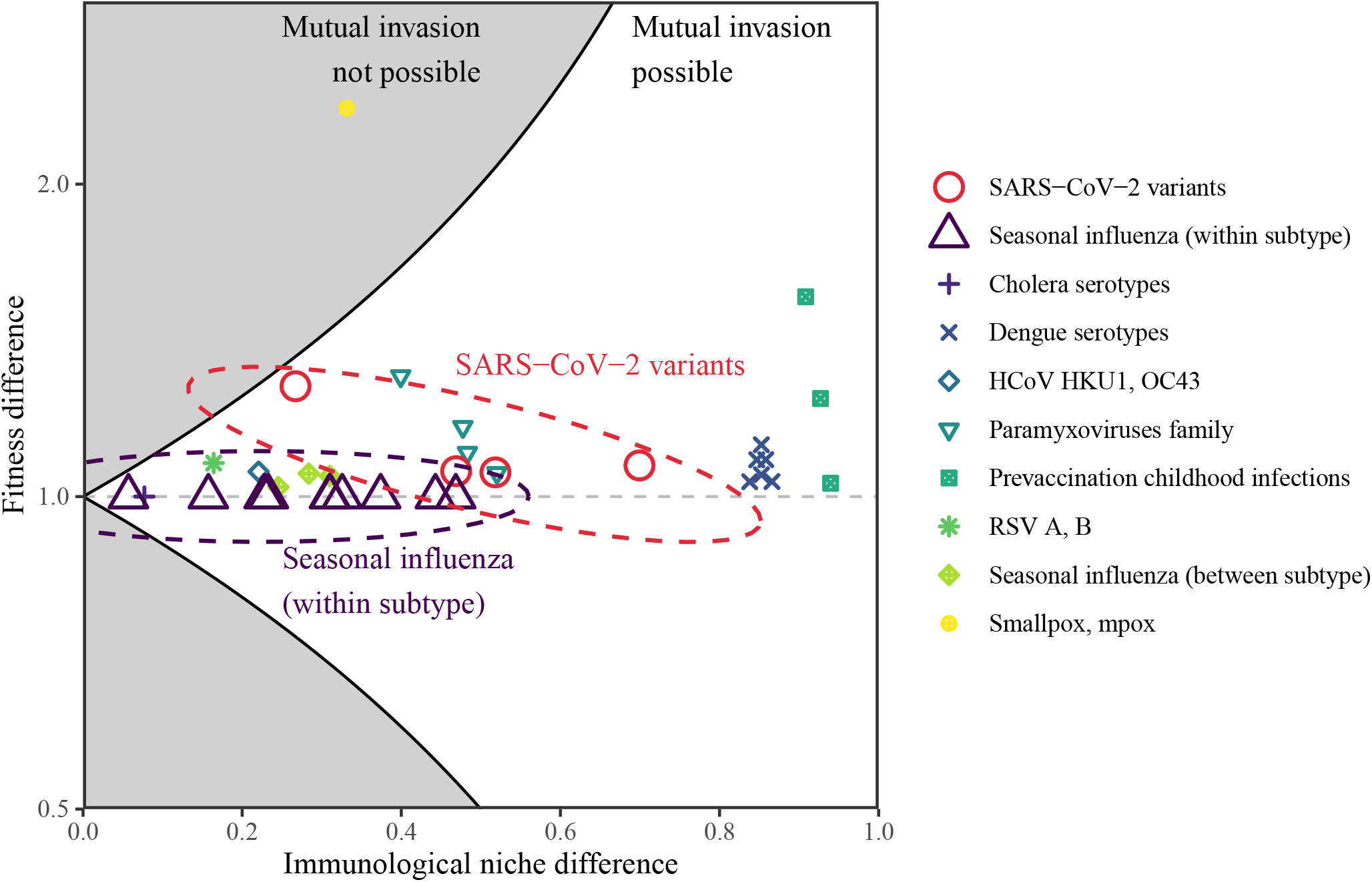
Quantifying coexistence patterns of different human pathogen variant pairs. Niche and fitness difference estimates across different human pathogen variants in the absence of vaccination. Each point indicates a pairwise comparison of pathogen variants from various systems, indicated by the shapes. Comparisons for prevaccination childhood infections considers ecological interference between measles vs chickenpox, rubella, and whooping cough (*1*). Points in the white region allow mutual invasion, meaning that a variant can invade a population when the competing variant is at equilibrium. Points in the grey region do not allow mutual invasion, meaning that the presence of more fit variant prevents the invasion of less fit variant. Dashed ellipses indicate SARS-CoV-2 variant pairs and within-subtype seasonal influenza strain pairs, which exhibit strain replacement patterns. We plot max(*f*_*i*_*/f*_*j*_, *f*_*j*_*/f*_*i*_) such that fitness difference is always greater than 1. The fitness difference is plotted on a log scale for symmetry.

For nearly all pathogen variant pairs tested, PIT predicts mutual invasion (Fig. 2). Two main factors contribute to this near universality of mutual invasibility. First, infection with one variant almost always confers weaker heterotypic immunity against other related pathogens, or their respective variants, than homotypic immunity; this in turn creates an immunological niche difference. Second, most closely related pathogens or pathogen variants frequently have similar *ℛ*_0_, which causes limited variation in the resulting fitness difference. Importantly, these two factors favoring mutual invasion also apply to antigenically variable strains of seasonal influenza for which strain replacement is the norm (*8, 30–32*). Similarly, even though SARS-CoV-2 variants are thought to have larger differences in their transmissibility (*33, 34*), we predict that their immunological niche difference is usually sufficiently large to permit mutual invasion (Fig. 2). Accounting for the effects of vaccination (Figure S1) or assuming a greater transmission advantage (Figure S2) prevents the SARS-CoV-2 WT variant from invading the SARS-CoV-2 Alpha variant; however, mutual invasibility predictions for all other variants are generally robust across different assumptions about vaccination, increased transmissibility, and cross immunity (Section S8; Figure S1–S3). One exception is smallpox in competition with mpox; in this case, PIT predicts that the endemic presence of smallpox would prevent the invasion of mpox. This prediction is consistent with previous hypotheses that mpox invasion was only possible because of smallpox eradication and the eventual waning of immunity against smallpox (*2, 28*). The apparent contradiction between the predicted mutual invasibility of seasonal influenza strains and SARS-CoV-2 variants (Fig. 2) and the observed patterns of competitive replacement (Fig. 1) raises a critical question of what mechanisms determine the co-circulation versus replacement of variants that can mutually invade one another.

### Mutual invasion, persistence, and co-circulation

The fate of pathogen competitors following invasion is determined by overcompensatory susceptible depletion dynamics, which are inherent to host-pathogen systems with significant acquired transmission-blocking immunity (*35*): an epidemic overshoots its peak, driving susceptibles below an invasion threshold. This overcompensation can prevent either or both competitors from persisting through an epidemic trough until the susceptible pool is replenished (*8, 9, 36*). In other words, co-circulation requires both resident and invading variants to overcome this overcompensatory effect.

For example, a classic RSV model (*25*) predicts that the invasion of either RSV A or B will cause both competitors to quickly settle to their endemic cycles, which match the observed dynamics (Fig. 3A). This contrasts with the strain replacement observed and modeled for SARS-CoV-2 and influenza (Fig. 1D,E). What factors allow RSV, but not SARS-CoV-2 or influenza, to overcome these overcompensatory effects? In order to synthesize the different competitive outcomes across these three major pathogen systems, we explore how subtle changes in immuno-epidemiological factors affect overcompensatory dynamics using the same

**Figure 3.**
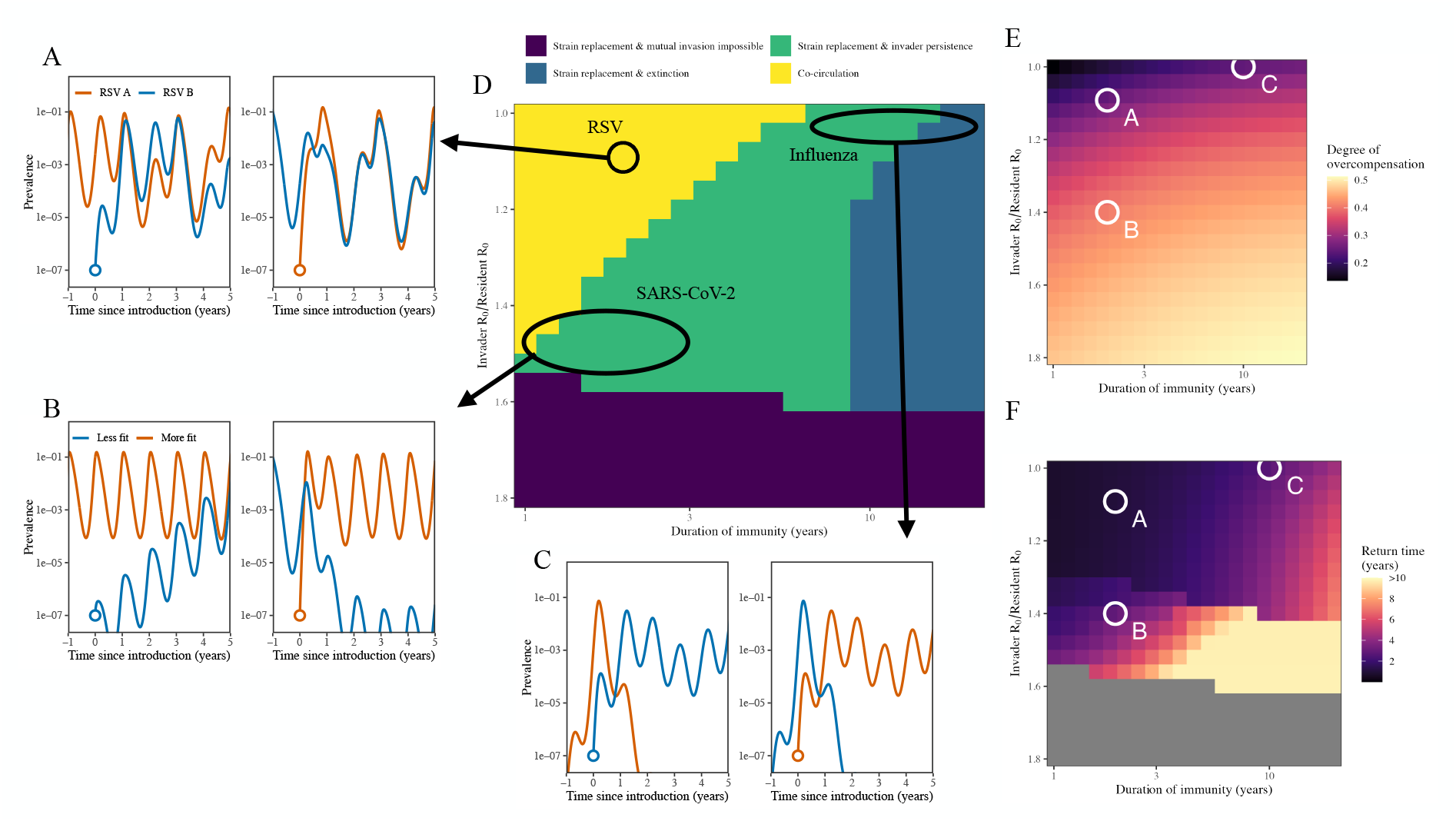
Effects of immunological factors on overcompensatory dynamics and competitive outcomes. (A–C) Example mutual invasion simulations that capture competitive outcomes of (A) RSV subtypes, (B) SARS-CoV-2 variants, and (C) seasonal influenza strains within subtype. Simulations assume (A) the same parameter estimates by (*25*); (B) 40% higher ℛ_0_ for the invading variant; and (C) equal ℛ_0_ for both variants and 10 years for the average duration of immunity. All other parameters are identical to the estimates by (*25*). (D) Competitive outcomes based on 10 years of simulations following the invasion of a novel variant while the resident is at equilibrium. Mutual invasion is possible in yellow, green, and blue regions. Mutual invasion is not possible only in the purple region. Co-circulation implies persistence of both resident and invading variants for 10 years. Persistence is measured by a prevalence cutoff of 10^−7^. All other parameters were fixed to previously estimated values. (E–F) Estimates of the degree of overcompensation and return time for the resident variant that correspond to simulations in panel D. For illustrative purposes, seasonal transmission was not included in for simulations in panels D–F. The standard RSV model by (*25*) was used for all simulations.

RSV model (Section S5). Specifically, we vary the duration of immunity and relative ℛ_0_ of the invading variant and characterize changes in competitive outcome (Fig. 3D), degree of over-compensation (Fig. 3E), and return time (Fig. 3F). The degree of overcompensation, defined as the maximum reduction in ℛ (*t*) (i.e., 1 − min(ℛ (*t*))) following the invasion of a novel variant, measures the depth of the trough. The return time, defined as the time between when ℛ (*t*) first decreases below 1 and ℛ (*t*) first increases above 1 again, measures the duration of the trough. Using ℛ (*t*) allows us to measure changes in the susceptible pool in relation to ℛ_0_.

We begin with the conditions that generate the co-circulation of mutually invading pathogens. Under short-term immunity and a limited transmission advantage for the invading variant (cor-responding to RSV), the model predicts prolonged co-circulation without local extinction of either variant (Fig. 3D, yellow region). This prediction is explained by the limited overcompensation (Fig. 3E) and fast return time of ℛ (*t*) to 1 (Fig. 3F). This dynamical regime also includes pneumococcal serotype competition (*11*), where the underlying dynamics exhibit minimal overcompensation and are therefore logistic-like, similar to standard ecological competitive systems (*23*). In this case, PIT further allows us to quantify how different immune mechanisms contribute to the maintenance of diversity (Sections S6, S9; Figure S4). In what follows we show that the replacement of mutually invading strains can be caused by either severe overcompensation or modest overcompensation coupled with a slow return time.

When the transmission advantage of an invading variant is large, the invasion of a more transmissible variant can cause strong overcompensation (Fig. 3E) and slower return times (Fig. 3F) even when the duration of immunity is short. This effect can prevent the resident variant from recovering and causes strain replacement despite mutual invasion (Fig. 3D, green region). This prediction suggests that the increased transmissibility of successive SARS-CoV-2 variants (*33, 34, 37*) was likely the major drivers of their replacement (Fig. 3B). This prediction is further supported by invasion simulations from a realistic SARS-CoV-2 model, which shows that the increased transmissibility is necessary for replacement (Section S10; Figures S5–S7). Here, mutual invasibility implies that the less transmissible variant can also invade the more transmissible variant, which would then lead to co-circulation (Fig. 3B). However, when the transmission advantage is too large, mutual invasion is not possible (Fig. 3D, purple region) and the more transmissible variant excludes the less transmissible variant.

Finally, even though longer-term immunity has limited impact on the degree of overcompensation (Fig. 3E), it can lead to a slow return time (Fig. 3F), explaining the replacement among seasonal influenza strains within a subtype (Fig. 3C,D). In fact, when the duration of immunity is very long, overcompensatory dynamics can cause local extinction for both invader and resident variants after strain replacement (Fig. 3D, blue region). These observations are further supported by more detailed influenza simulations, which show that prolonged immunity against individual seasonal influenza strains of the same subtype (*38–40*) makes co-circulation rare despite near-universal mutual invasibility (Sections S7, S11; Figures S8–S10).

### Summary

There has been extensive work analyzing variant invasion and coexistence across many individual host-pathogen systems; however, synthesis and comparison across systems is lacking. Here, we present a unifying framework, PIT, that allows for such comparison by quantifying key coexistence metrics—niche and fitness differences—in two-pathogen systems via the analysis of mechanistic models (Fig. 4). PIT reveals that mutual invasion of initially rare variants is nearly universal for major pathogen systems, including for exemplary systems of strain replacement such as seasonal influenza and SARS-CoV-2. Mutual invasibility equates to stable coexistence in ecological systems dominated by logistic Lotka-Volterra-like dynamics (*23*).

**Figure 4.**
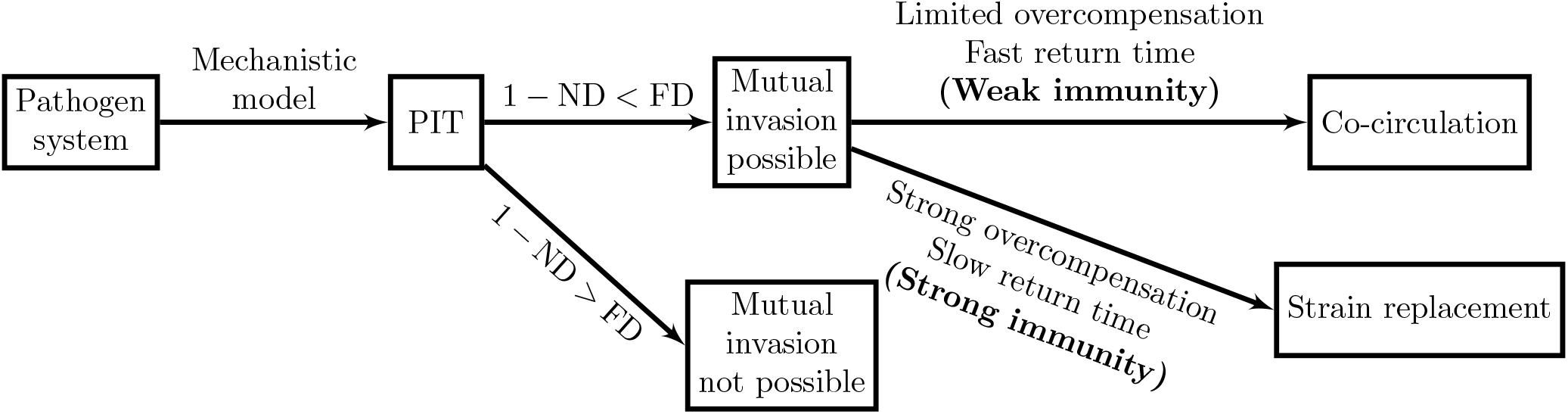
Schematic summary of PIT and predictions for long-term co-circulation. Pathogen invasion theory (PIT) allows predictions of mutual invasibility based on estimates of the immunological niche difference (ND) and fitness difference (FD). These estimates require well-parameterized mechanistic models for a given two-pathogen system. Even when two variants can mutually invade, predictions for strain replacement depend on the degree of overcompensation and return time from susceptible depletion.

By contrast, whether the mutual invasion of pathogen variants leads to co-circulation or replacement is determined by the extent of overcompensatory dynamics of susceptible (resource) depletion (*9, 36, 41*). Specifically, the degree of overcompensation and return times determine whether a variant can persist. For example, when both invading and resident variants have similar transmissibility, the degree of overcompensation is limited. In this case, the duration of immunity dictates return time from susceptible depletion and therefore is the key determinant for pathogen co-circulation: short-term immunity allows co-circulation (e.g., between RSV sub-types), but prolonged immunity leads to replacement (e.g., between seasonal influenza strains within the same subtype). Increasing the transmission advantage of an invading variant leads to stronger overcompensation, making pathogen persistence more difficult. In this case, strong overcompensation can lead to replacement even under short-term immunity (e.g., SARS-CoV-2 variants).

### Caveats

A key limitation of PIT is that it focuses on pairwise comparisons. This is in contrast to the observed competition patterns (Fig. 1), which may depend on multi-pathogen interactions as well as the longer-term, multi-pathogen immune history. The importance of community structure and higher-order interactions, which only arise in the presence of more than two competing species, is a major unknown in community ecology (*42, 43*). Similar effects of higher-order interactions can be found in host-pathogen systems: for example, infection with two different dengue serotypes can provide strong immunity against subsequent infections against all other serotypes (*44*). We also focus on analyzing major, endemic pathogen systems through PIT, but there are many rare pathogen systems (*7*); nonetheless, we generally expect conclusions about near-universality of mutual invasibility to be robust given that they hold for exemplary systems of replacement (i.e., seasonal influenza and SARS-CoV-2). Finally, PIT focuses on invasion at the endemic attractor, but the dynamics of many pathogens, especially those that exhibit strain replacement patterns, can be far from the attractor.

Our invasion analyses focus on competition within single, well-mixed host populations and therefore neglect spatial variation. This likely renders our predictions about pathogen coexistence conservative. In reality, spatial variation and metapopulation structure can play an important role in determining overcompensation and maintaining pathogen persistence at regional scales (*45*). Seasonality can also affect overcompensation and contribute to pathogen persistence (*46*). Despite these limitations, our work provides a quantitative foundation for understanding pathogen coexistence.

## Conclusion

The recent emergence and re-emergence of novel pathogens has caused significant disruption to our society, necessitating better surveillance and control strategies to prevent future outbreaks. The near universality of pathogen mutual invasibility raises a serious public health concern given the potential recombination among variants and the emergence of new pandemic variants (*47*). Our results further underline the importance of global pathogen and immunological surveillance platforms for clarifying the dynamics of a potentially diverse range of future threats (*48*). A detailed understanding of immuno-epidemiological dynamics will be critical to predicting the persistence and community structure of novel pathogens in the future.

## Acknowledgments

We thank Jonathan Dushoff for helpful discussion and comments on early versions of this manuscript.

## Funding

S.W.P. was supported by Charlotte Elizabeth Procter Fellowship of Princeton University. C.J.E.M. and B.T.G were supported by Princeton Catalysis Initiative and Princeton Precision Health. J.M.L was supported by the National Science Foundation (2022213). S.C. was supported by Federal funds from the National Institute of Allergy and Infectious Diseases, National Institutes of Health, Department of Health and Human Services under CEIRR contract 75N93021C00015. The content is solely the responsibility of the authors and does not necessarily represent the official views of the NIAID or the National Institutes of Health.

## Author contributions

S.W.P. conceived of the study, conducted the analysis, and wrote the manuscript. S.C. assisted with the analysis and edited the manuscript. C.J.E.M. conceived of the study, assisted with the analysis, edited the manuscript, and oversaw the work. J.M.L. conceived of the study, assisted with the analysis, wrote the manuscript, and oversaw the work. B.T.G. conceived of the study, assisted with the analysis, wrote the manuscript, and oversaw the work.

## Competing interests

The authors declare no competing interests.

## Data availability

All data used in this paper are publicly available in references provided in Fig. 1. All code used in this paper are available on https://github.com/parksw3/invasion.

## Supplementary materials

Materials and Methods

Supplementary Text

Figs. S1 to S10

References (49–58)

## Supplementary Materials for

### Materials and Methods

#### S1 Chesson’s modern coexistence theory

Chesson’s modern coexistence theory (MCT) stems from a Lotka-Volterra system, where the population dynamics of species *i* can be written as

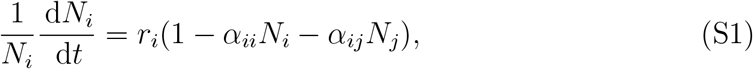

where *r*_*i*_ represents the growth rate of species *i, α*_*ii*_ represents the strength of intraspecific competition, and *α*_*ij*_ represents the strength of interspecific competition. Then, mutual invasibility requires a stronger intraspecific competition than interspecific competition: *α*_*ii*_ *> α*_*ji*_ (and *α*_*jj*_ *> α*_*ij*_). Alternatively, these conditions can be also expressed in terms of niche overlap *ρ* and average fitness difference *f*_*i*_*/f*_*j*_ (*23*):

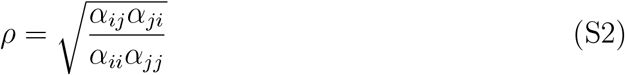

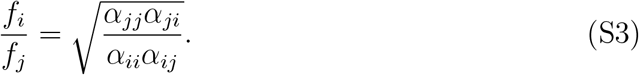

#### S2 Pathogen invasion theory

For any two pathogen variants *i* and *j*, their equilibrium states must satisfy (*35*)

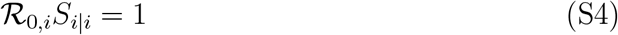

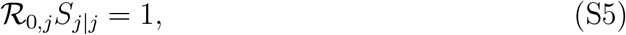

where ℛ_0,*i*_ represents the basic reproduction number of variant *i* and *S*_*i*|*i*_ represents the effective proportion of individuals who are susceptible to variant *i* when variant *i* is at equilibrium (and likewise for ℛ_0,*j*_ and *S*_*j*|*j*_). For mutual invasion, an invading variant, initially rare, has to be able to invade the competing variant at its equilibrium; these conditions can be written as

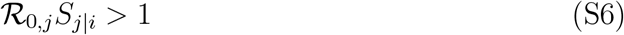

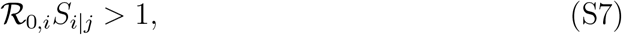

where *S*_*j*|*i*_ represents the effective proportion of individuals who are susceptible against variant *j* when variant *i* is at equilibrium (and vice versa for *S*_*i*|*j*_). Alternatively, the mutual invasibility criteria can be rewritten as

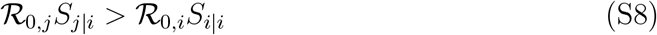

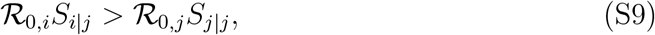

which suggests that variant *i* must reduce its own transmission ℛ_0,*i*_ more than it reduces the transmission of a competing variant ℛ_0,*j*_—this observation strongly echoes a key foundation in MCT that intraspecific competition must be stronger than interspecific competition. This mutual invasibility criteria can be further rewritten as

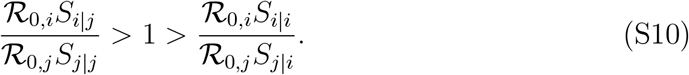

Dividing both sides by the fitness difference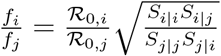, we get

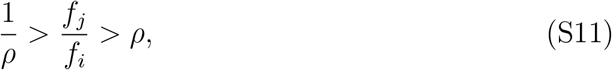

where 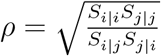 represents the niche overlap. We note that the fitness difference can be equivalently written as

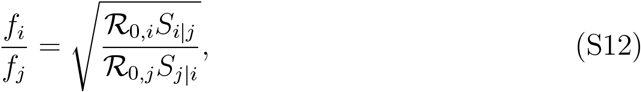

which simply measures the ratio of how well each variant spreads when the competing variant is at equilibrium.

#### S3 Effects of pathogen interactions on susceptibility, transmissibility, and duration of infection

The original definitions of niche overlap and fitness difference in Section S2 only account for the effects of pathogen interactions on population-level susceptibility. However, pathogen interactions, especially cross immunity, can also affect transmissibility as well as duration of infections. In order to account for changes in susceptibility, transmissibility, and duration of infection simultaneously, we begin by writing *S*_*i*|*h*_(*t*) to denote the proportion of individuals who are susceptible to variant *i* at time *t* given history *h*, where *h* can depend on both previous history of infection or current carriages of the same or different variants. Likewise, we write *I*_*i*|*h*_(*t*) are infected with variant *i* at time *t* given history *h* such that infections of individuals in the *S*_*i*|*h*_(*t*) compartment will cause them to move to the *I*_*i*|*h*_(*t*) compartment. Then, we have

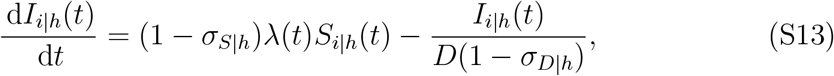

where *λ*(*t*) represents the force of infection at time *t, σ*_*S*|*h*_ represents the proportional reduction in susceptibility given history *h, D* represents the mean duration of infection of a naive individual, and *σ*_*D*|*h*_ represents the proportional reduction in the duration of infection given history *h*. Assuming a homogeneously mixing population, the force of infection can be calculated by adding up transmission from all infection-history classes:

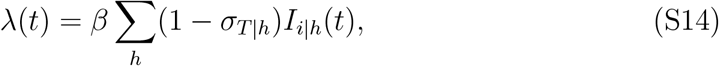

where *σ*_*T* |*h*_ represents the proportional reduction in transmission rate given history *h*.

Endemic equilibrium is described by the following condition (across all *h*):

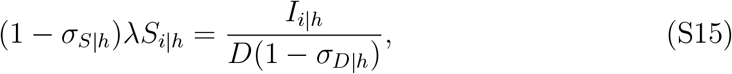

which can be re-written as

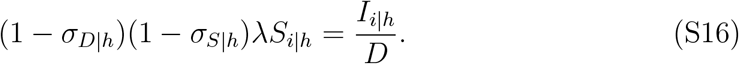

Then, by substituting the above expression for *λ* and summing across all possible history *h*, we get

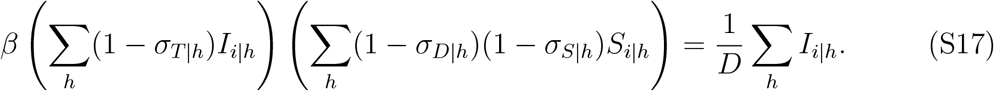

By moving everything on the right-hand side to the left-hand side, we get

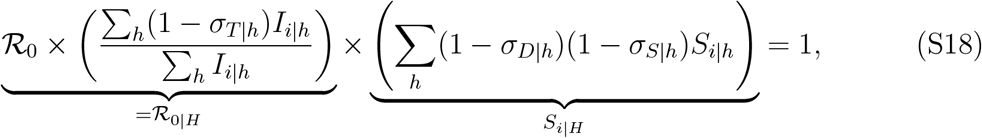

where the first term, ℛ_0_ = *βD*, represents the basic reproduction number, which is defined as the average number of secondary infections caused by an infected individual in a fully susceptible population; the second term (∑_*h*_(1 − *σ*_*T* | *h*_)I_*i*|*h*_)*/*( ∑_*h*_ *I*_*i*|*h*_) captures average proportional reduction in transmission rate at equilibrium; and the remaining term captures the average proportional reduction in the duration of infection and susceptibility at equilibrium. We note that the entire expression on the left hand side (i.e., the product of three terms) also corresponds to the effective reproduction number. We can then rewrite the whole expression as

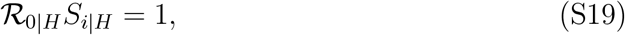

wheℛ*R*_0|*H*_ represents the average reproduction number across the set of all possible infection history *H* (accounting for the impact of immunity on transmission reduction), and *S*_*i*|*H*_ represents the mean susceptibility across the set of all possible infection history *H*(accounting for the impact of immunity on the reduction of susceptibility and infection duration).

Now, deriving measures for niche overlap and fitness differences is straightforward. At equilibrium, we have

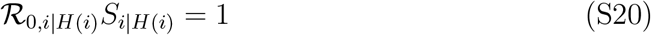

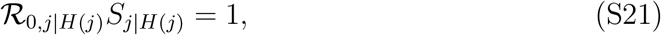

where *H*(*i*) and *H*(*j*) denotes the set of all possible infection history when variant *I* and *j* are at equilibrium, respectively. For mutual invasion, we need

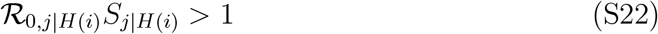

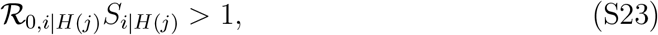

which captures the average effect on variant *j* of all infection history when variant *i* is at equilibrium (and vice versa). Note that calculating ℛ_0,*j*|*H*(*i*)_ (and likewise, ℛ_0,*i*|*H*(*j*)_) depends on the distribution of infection prevalence *I*_*i*|*h*(*j*)_ (and *I*_*j*|*h*(*i*)_) during the growth phase, which is determined by the eigenvector, rather than at the equilibrium. Then, mutual invasibility criteria can be written as

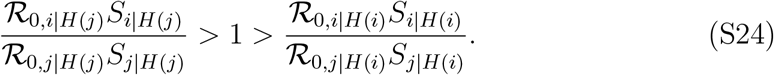

In this case, the fitness difference is given by:

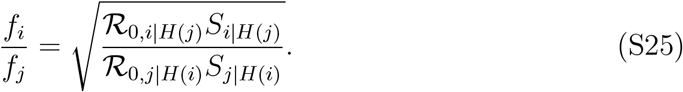

Dividing both sides of the mutual invasibility criteria gives us:

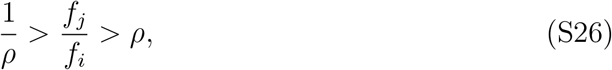

where the niche overlap *ρ* corresponds to:

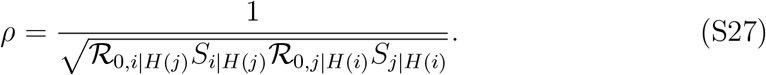

This expression generalizes the expression presented in Section S2: when ℛ_0,*i*|*H*(*j*)_ = ℛ_0,*i*|*H*(*i*)_ and ℛ_0,*j*|*H*(*i*)_ = ℛ_0,*j*|*H*(*j*)_ (i.e., when intraspecific and interspecific effects of transmission reduction are identical), we get the same expression presented in Section S2.

#### S4 Quantifying niche and fitness differences for the observed pathogen variants

We collated a series of mechanistic models that were previously directly fitted to pathogen time series data, or indirectly parameterized based on data, and computed niche and fitness differences either analytically or numerically using simulations (*19, 1, 24, 25, 26, 27, 11, 28, 14, 29, 15*). Here, we present all models that we used and corresponding simulation setups and parameter sets.

##### S4.1 Cholera model

Cholera dynamics are modeled based on a two-strain SIR model, which assumes that infection with one serotype provides permanent immunity against the same serotype (*26*). The model also assumes that current or past infection by one serotype reduces susceptibility against another serotype by *σ*. Then, mean susceptibility at endemic equilibrium satisfies the following:

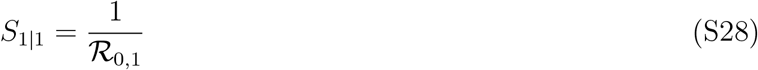

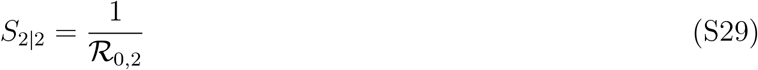

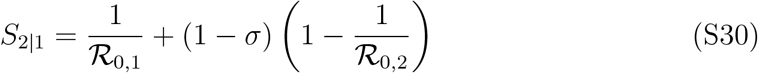

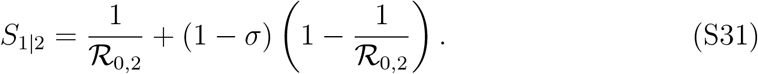

Following (*26*), we assume *σ* = 0.956, ℛ_0,1_ = ℛ_0,2_ = 2.9 to calculate niche and fitness difference. This parameterization qualitatively captures complex serotype cycles observed in Bangladesh when stochasticity is accounted for. We note that the same analytical expressions can be used for the SIRS model, and therefore, this simple approach can be used for pathogens that permit reinfection.

##### S4.2 Seasonal human coronavirus model

Seasonal human coronavirus dynamics are modeled based on a two-strain SEIRS model, which includes the exposed stage, during which individuals are infected but not yet infectious, and allows immunity to wane (*14*). This model was previously fitted to HCoV-OC43 and HCoV-HKU1 time series from the US. The dynamics are described by the following set of ordinary differential equations (see Figure S4 of (*14*) for the model diagram):

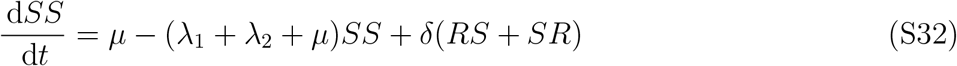

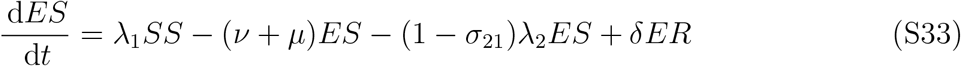

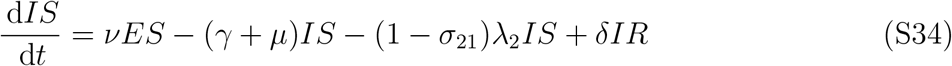

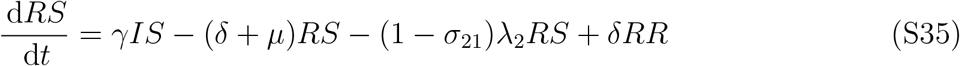

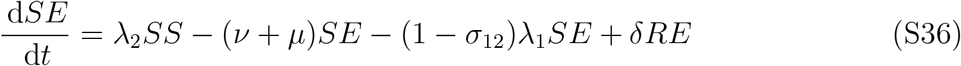

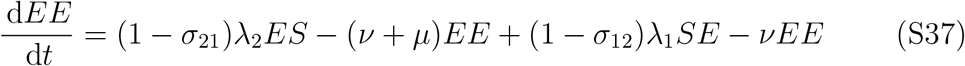

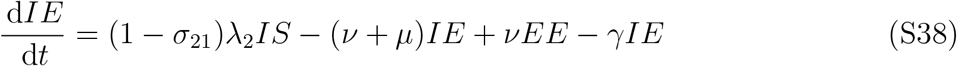

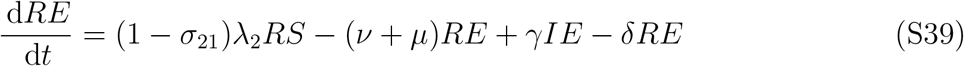

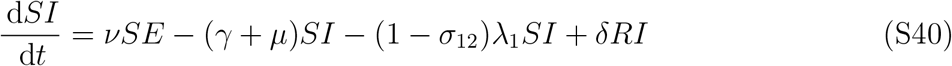

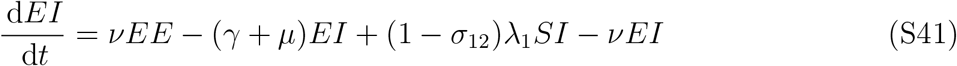

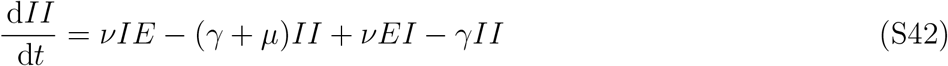

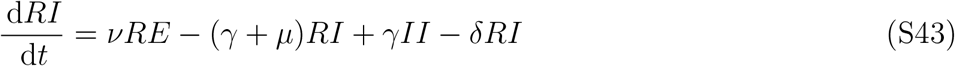

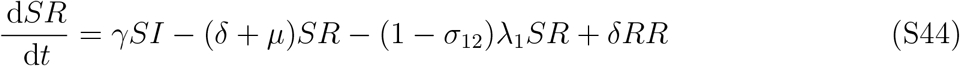

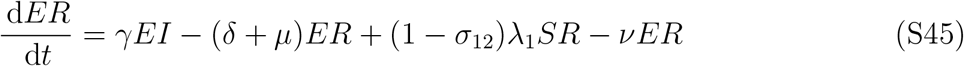

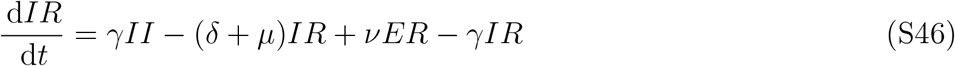

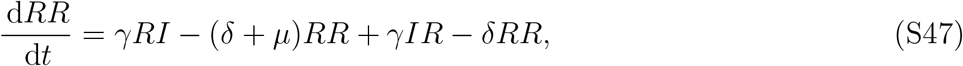

where each compartment *XY* describes the infection status against the first (*X*) and second (*Y*) strains; *λ*_*i*_ represents the force of infection of strain *i*; *µ* represents birth/death rates; *δ* represents the rate of waning immunity; *ν* represents the rate at which exposed individuals become infectious; *γ* represents the rate of recovery; and *σ*_*ij*_ represents the strength of cross immunity of strain *j* against strain *i*. Here, the force of infection is defined as follows:

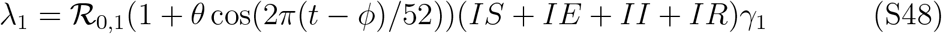

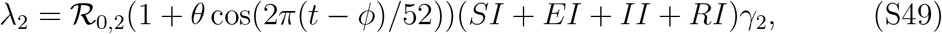

where *θ* represents the seasonal amplitude; and *ϕ* represents the offset term.

For this model, mean susceptibility at endemic equilibrium corresponds to:

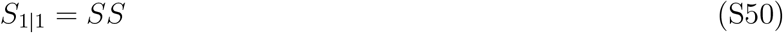

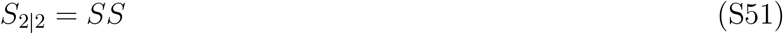

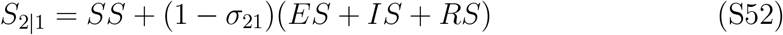

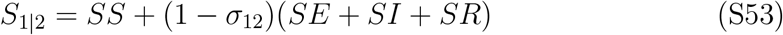

which are calculated by introducing one variant at a time and running the simulation for 100 years, allowing the system to reach its endemic attractor. Niche and fitness difference measures are calculated by taking the long-term average across last 10 years of simulations. We assume following parameter values based on (*14*): 1*/ν* = 5*/*7 weeks, 1*/γ* = 5*/*7 weeks, *σ*_21_ = 0.7, *σ*_12_ = 0.5, ℛ_0,1_ = ℛ_0,2_ = 1.7, *θ* = 0.3, *ϕ* = −4, 1/δ = 40 weeks, and 1*/µ* = 52 *×* 80 weeks.

##### S4.3 Between-subtype seasonal influenza model

Between-subtype influenza dynamics are modeled based on the model used by (*29*), which extends the model by (*30*) to include waning immunity. The model was previously fitted to seasonal influenza time series from Hong Kong. The dynamics are described by the following set of differential equations:

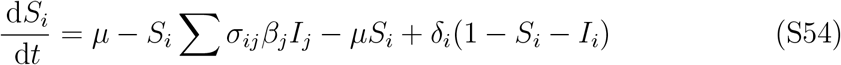

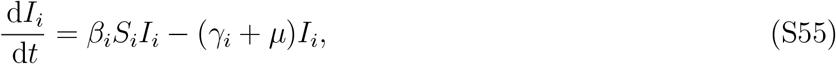

where *S*_*i*_ and *I*_*i*_ represent the proportion of individuals who are susceptible to and infected with strain *i*, respectively; *σ*_*ij*_ represents the strength of cross immunity of strain *j* against strain *i*; *µ* represents the birth rate; *δ* represents the rate of waning immunity; *γ* represents the rate of recovery; and *β* represents the transmission rate. The transmission rate is modeled using a sinusoidal function: *β*_*i*_ = ℛ_0,*i*_(1 + *θ* cos(2*πt/*52))(*µ* + *γ*_*i*_), where *θ* represents the seasonal amplitude.

For this model, mean susceptibility at endemic equilibrium is calculated by introducing one strain and taking *S*_*i*_ for the corresponding strain at equilibrium. For example, we can simply take *S*_*i*|*i*_ = *S*_*i*_ and *S*_*j*|*i*_ = *S*_*j*_ after introducing the *i*-th strain. To calculate niche overlap and fitness difference measures, we run the simulation for 200 years and take the long-term average across last 20 years of simulations. We assume following parameter values based on (*29*): ℛ_0,1_ = 1.44, 1*/γ* = 2.64*/*7 weeks, 1*/δ* = 3.12*×*52 weeks for A/(H1N1); ℛ_0,1_ = 1.60, 1*/γ* = 3.03*/*7 weeks, 1*/δ* = 2.28*×*52 weeks for A/(H3N2); ℛ_0,1_ = 1.43, 1*/γ* = 3.09*/*7 weeks, 1*/δ* = 3.08 *×* 52 weeks for B; and 1*/µ* = 52 *×* 80 weeks across all simulations. The original model tried to estimate separate transmission rates for each week. Instead, we use a sinusoidal function with *θ* = 0.1 in order to capture the dynamics of the model at its endemic attractor.

##### S4.4 RSV model

RSV dynamics are modeled based on the classic model used by (*25*). The model was previously fitted to RSV time series data from England & Wales and Finland (see Figure 1 of the original paper for a model diagram).

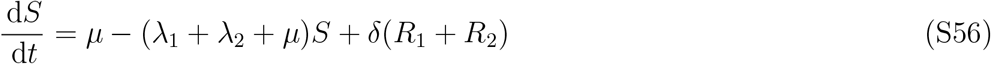

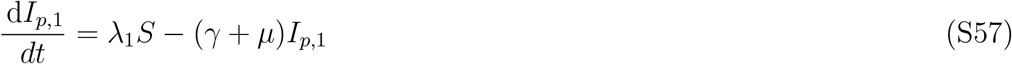

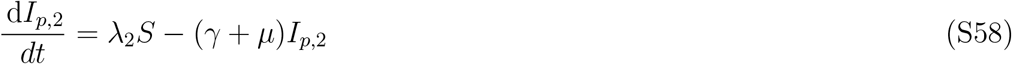

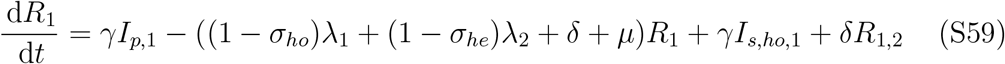

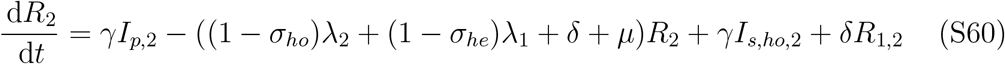

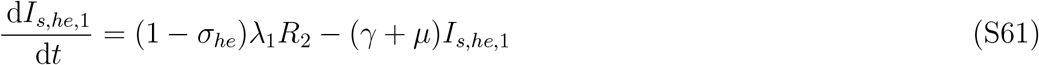

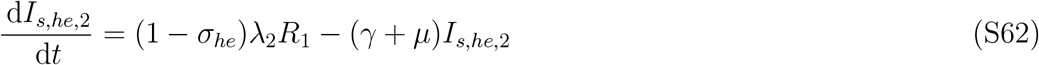

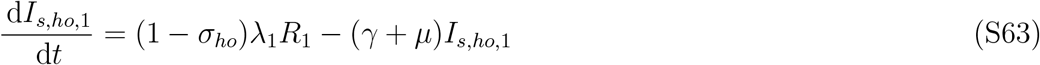

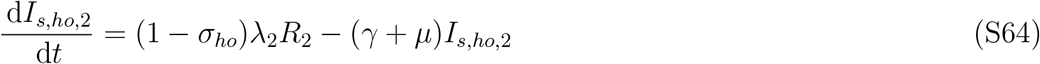

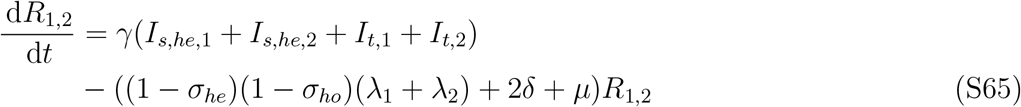

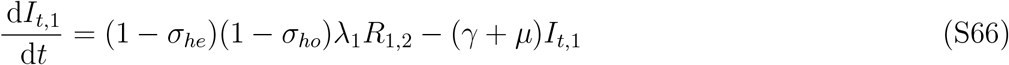

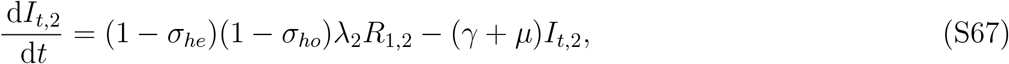

where *S* represents the proportion of individuals who are fully susceptible to both infections; *I*_*p*,1_ and *I*_*p*,2_ represent the proportion of individuals who have primary infection; *R*_1_ and *R*_2_ represent the proportion of individuals who have recovered from primary infection; *I*_*s,he*,1_ and *I*_*s,he*,2_ represent the proportion of individuals who have heterotypic secondary infection; *I*_*s,ho*,1_ and *I*_*s,ho*,2_ represent the proportion of individuals who have homotypic secondary infection; *R*_1,2_ represents the proportion of individuals who have recovered from secondary or tertiary infections; *I*_*t*,1_ and *I*_*t*,2_ represent the proportion of individuals who have tertiary infections; *µ* represents the birth/death rate; *λ* represents the force of infection; *δ* represents the rate of waning immunity; *γ* represents the rate of recovery; *σ*_*ho*_ represents the strength of homotypic immunity; and *σ*_*he*_ represents the strength of heterotypic immunity. The force of infection corresponds to

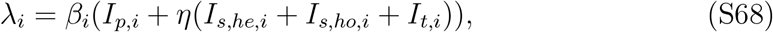

where *β* represents the transmission rate; and *η* represents reduction in transmission rate for reinfection classes. The transmission rate is assumed to follow a sinusoidal function:

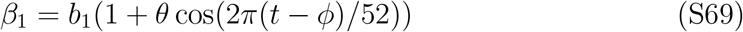

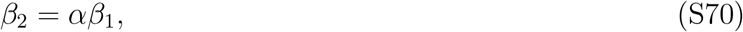

where *b*_1_ represents the mean transmission rate of strain 1; *θ* represents the seasonal amplitude; *ϕ* represents the offset term; and *α* represents the relative transmission rate of RSV B compared to RSV A.

For this model, mean susceptibility at endemic equilibrium corresponds to

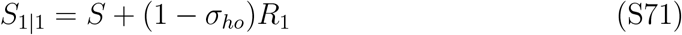

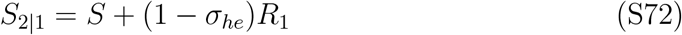

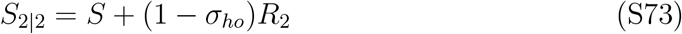

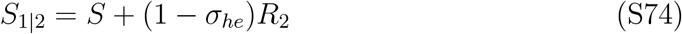

Since only one RSV subtype is present at endemic equilibrium, we do not have to consider tertiary infections for PIT calculations.

This model also includes the impact of immunity on transmission reduction, so we follow the approach outlined in Section S3 to account for reduction in transmission rate. Specifically, we calculate

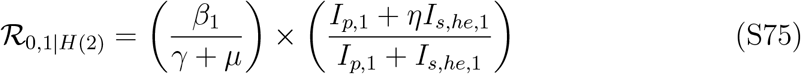

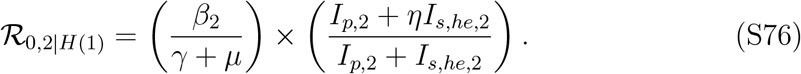

We note that ℛ_0,1|*H*(2)_ and ℛ_0,2|*H*(1)_ need to be calculated during the exponential growth phase of an invasion of the invader while the resident is at its equilibrium. To do so, we first run the model with a resident strain only for 100 years and then introduce the invading strain at an extremely low prevalence (10^−100^) and running it for 10 more years. We also note that the term *I*_*t*,1_ and *I*_*t*,2_ can be ignored for this calculation because they are new infections arising from *R*_1,2_. Since *R*_1,2_ requires infection from both subtypes, the proportion of individuals in this compartment is equal to zero when only one variant is at equilibrium. Likewise, the contribution from this compartment is expected to be negligible during the initial invasion. Mean susceptibility at endemic equilibrium is calculated from the first set of simulations (i.e., using the final 10 years of the 100 year simulation). We assume following parameter values based on parameter estimates for Finland (*25*): *α* = 0.9159, *η* = 0.4126, *σ*_*ho*_ = 0.6431, *σ*_*he*_ = 0.1574, *γ* = 40.56*/*52*/*year, *δ* = 0.51*/*52*/*year, *b*_1_ = 99.51*/*52*/*year, *µ* = 0.012*/*52*/*year, *ϕ* = 0.97 *×* 52, and *θ* = 0.347.

##### S4.5 Paramyxovirus family model

Epidemic dynamics of paramyxovirus family are modeled based on the model used by (*13*). This model was previously fitted to time series data of infections by paramyxovirus family from Utah and Idaho. The model equations are given by:

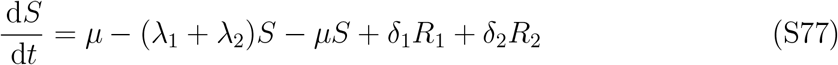

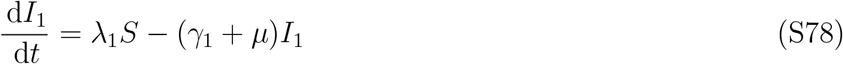

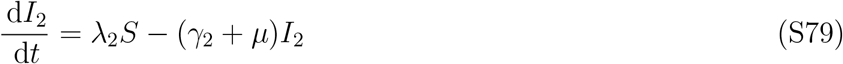

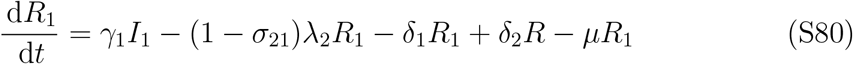

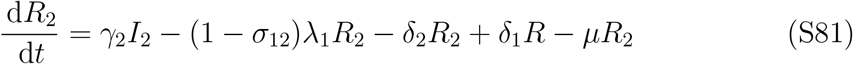

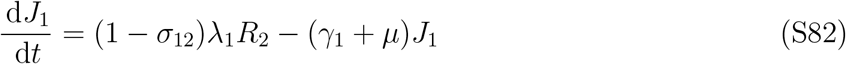

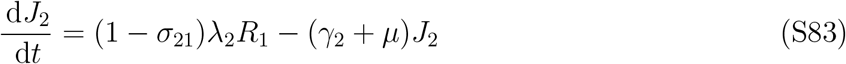

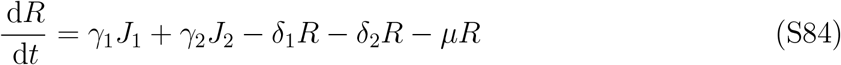

where *S* represents the proportion of individuals who are fully susceptible to both infections; *I*_1_ and *I*_2_ represent the proportion of individuals who have primary infection; *R*_1_ and *R*_2_ represent the proportion of individuals who have recovered from primary infection; *J*_1_ and *J*_2_ represent the proportion of individuals who have secondary infection; *µ* represents the birth/death rate; *λ* represents the force of infection; *δ* represents the rate of waning immunity; *γ* represents the recovery rate; and *σ*_*ij*_ represents the strength of cross immunity of pathogen *j* against pathogen *i*. The force of infection corresponds to

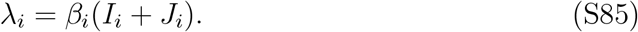

where *β* represent the transmission rate. The transmission rate is assumed to follow a sinusoidal function:

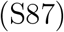

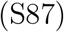

where *b*_*i*_ represents the mean transmission rate, *θ* represents the seasonal amplitude; *ϕ* represents the offset term; and *q* represents transmission probability.

For this model, mean susceptibility at endemic equilibrium corresponds to:

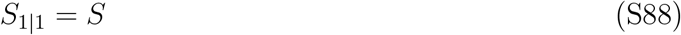

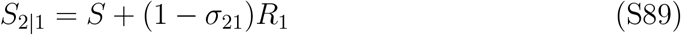

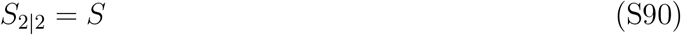

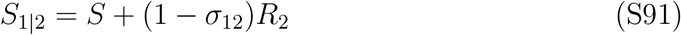

which are calculated by introducing one pathogen at a time and running the simulation for 30 years for it reach its endemic attractor. Niche and fitness difference measures are calculated by taking the long-term average across last 10 years of simulations. We assume following parameter values based on (*25*): *b*_1_ = 3.4*/*weeks, *b*_2_ = 2.9*/*weeks, *θ*_1_ = 0.4, *θ* = 0.33, *ϕ*_1_ = −17.5, ϕ_2_ = −29.5, σ_12_ = 0.1, and *σ*_21_ = 0.46 for the RSV (pathogen 1) and HPIV-1 (pathogen 2) pair; *b*_1_ = 3.4*/*weeks, *b*_2_ = 2.4*/*weeks, *θ*_1_ = 0.4, *θ*_2_ = 0.28, *ϕ*_1_ = −17.99, ϕ_2_ = −20.001, *σ*_12_ = 0.1, and *σ*_21_ = 0.204 for the RSV (pathogen 1) and HPIV-2 (pathogen 2) pair; *b*_1_ = 3.4*/*weeks, *b*_2_ = 2.0*/*weeks, *θ*_1_ = 0.4, *θ*_2_ = 0.31, *ϕ*_1_ = −18, ϕ_2_ = −11, σ_12_ = 0.13, and *σ*_21_ = 0.37 for the RSV (pathogen 1) and HPIV-3 (pathogen 2) pair; and *b*_1_ = 3.4*/*weeks, *b*_2_ = 3.9*/*weeks, *θ*_1_ = 0.4, *θ*_2_ = 0.3, *ϕ*_1_ = 0.005, *ϕ*_2_ = 4.99, *σ*_12_ = 0.08, and *σ*_21_ = 0.55 for the RSV (pathogen 1) and hMPV (pathogen 2) pair. Remaining parameters are held constant across different pairs: *γ* = 7*/*10*/*weeks, *ρ* = 1*/*52*/*weeks, *µ* = 1*/*70*/*52*/*weeks, and *q* = 0.5.

##### S4.6 Dengue serotype model

Epidemic dynamics of dengue serotypes are modeled based on the model used by (*19*), which assumes an SIR model for single serotype dynamics. Therefore, the mutual invasibility criteria can be calculated analytically in the same way as the Cholera model (Section S4.1). To account for the effect of antibody-dependent enhancement, we assume *σ* = −0.5, which means that primary infection increases susceptibility against future infections by 50%. Basic reproduction number estimates for each serotype is taken from (*49*): 4.29, 5.12, 4.61, and 5.50 for DENV 1, 2, 3, and 4, respectively.

##### S4.7 Prevaccination childhood infections model

Epidemic dynamics of prevaccination childhood infections are modeled based on the model used by (*1*), which accounts for ecological interactions across different infections. The model assumes that individuals who recover from infections enter a convalescence period, during which they are temporarily removed from the susceptible pool. The dynamics are described by the following set of equations:

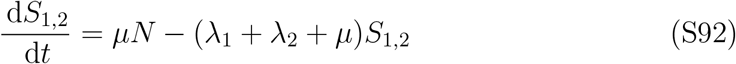

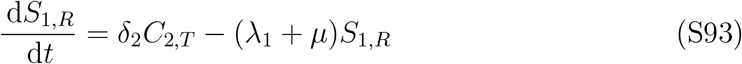

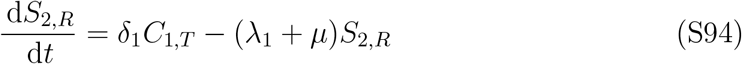

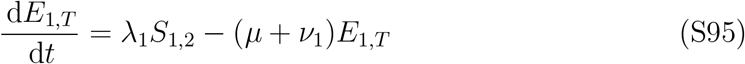

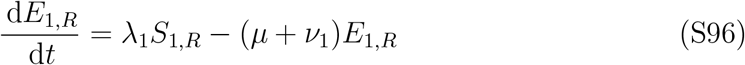

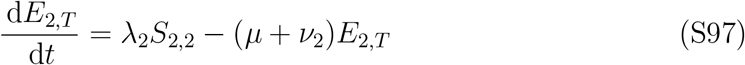

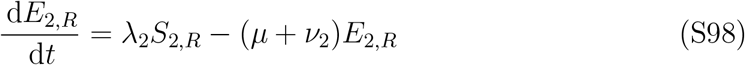

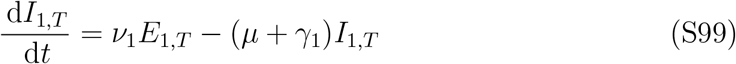

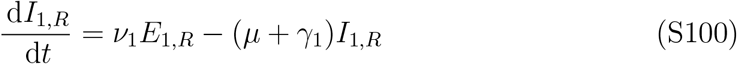

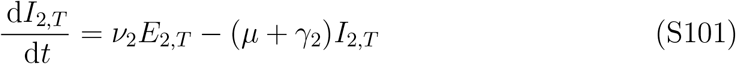

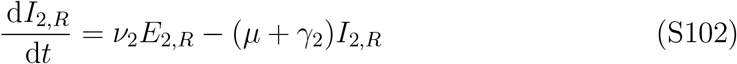

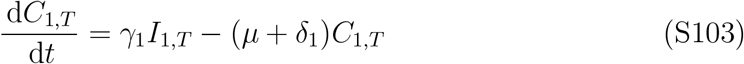

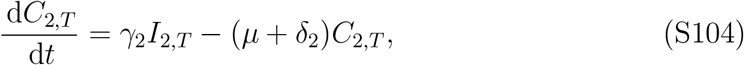

where *S*_1,2_ represents the proportion of individuals who are fully susceptible; *S*_1,*R*_ and *S*_2,*R*_ represent the proportion of individuals who have recovered from an infection with one pathogen but is still susceptible to the other pathogen; *E*_1,*T*_ and *E*_2,*T*_ represent the proportion of individuals who have been exposed to either pathogens for the first time; *E*_1,*R*_ and *E*_2,*R*_ represent the proportion of individuals who have recovered from an infection with one pathogen and have been exposed to a different pathogen; *I*_1,*T*_ and *I*_2,*T*_ represent the proportion of individuals who have been infected by either pathogens for the first time and are infectious; *I*_1,*R*_ and *I*_2,*R*_ represent the proportion of individuals who have recovered from an infection with one pathogen and have been exposed to a different pathogen and are infectious; *C*_1,*T*_ and *C*_2,*T*_ represent the proportion of individuals who have recovered from an infection with one pathogen and have been temporarily removed from the susceptible pool; *µ* represents the birth/death rate; *λ* represents the force of infection; *ν* represents the rate at which exposed individuals become infectious; *γ* represents the recovery rate; and *δ* represents the rate at which recovered individuals become susceptible to new infections.

The force of infection corresponds to

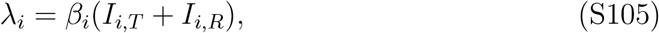

where *β* represents the transmission rate. The transmission rate is assumed to follow a sinusoidal function:

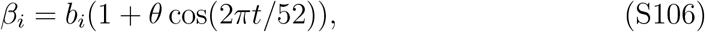

where *b*_*i*_ represents the mean transmission rate, and *θ* represents the seasonal amplitude.

For this model, mean susceptibility at endemic equilibrium corresponds to

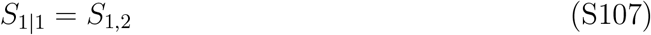

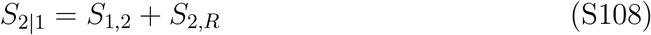

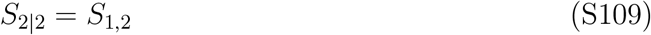

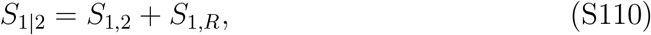

which are calculated by introducing one pathogen at a time and running the simulation for 200 years for it reach its endemic attractor. Niche and fitness difference measures are calculated by taking the long-term average across last 10 years of simulations. We assume following parameter values based on (*1*): *b*_1_ = 1250*/*52*/*weeks, *ν*_1_ = 7*/*8*/*weeks, *γ*_1_ = 7*/*5*/*weeks, *b*_2_ = 625*/*52*/*weeks, *ν*_2_ = 7*/*8*/*weeks, *γ*_2_ = 7*/*10*/*weeks, *δ*_1_ = 7*/*7*/*weeks, and *δ*_2_ = 7*/*14*/*weeks for measles and whooping cough pair; *b*_1_ = 1250*/*52*/*weeks, *ν*_1_ = 7*/*8*/*weeks, *γ*_1_ = 7*/*5*/*weeks, *b*_2_ = 49*/*11*/*weeks, *ν*_2_ = 7*/*9*/*weeks, *γ*_2_ = 7*/*11*/*weeks, *δ*_1_ = 7*/*7*/*weeks, and *δ*_2_ = 7*/*14*/*weeks for measles and rubella pair; and *b*_1_ = 1250*/*52*/*weeks, *ν*_1_ = 7*/*8*/*weeks, *γ*_1_ = 7*/*5*/*weeks, *b*_2_ = 77*/*10*/*weeks, *ν*_2_ = 7*/*10*/*weeks, *γ*_2_ = 7*/*10*/*weeks, *δ*_1_ = 7*/*7*/*weeks, and *δ*_2_ = 7*/*14*/*weeks for measles and chickenpox pair. Remaining parameters are held constant across all pairs: *µ* = 1*/*50*/*52*/*weeks, and *θ* = 0.2.

##### S4.8 Smallpox and mpox model

Niche and fitness difference between smallpox and mpox are calculated analytically based on the analytical approach shown in Section S4.1. Given large uncertainties in cross immunity between two pathogens as well as their transmissibility, we calculated niche and fitness differences across a wide range of realistic parameters and took their mean: ℛ_0,1_ = 3.5 − 6 for smallpox (*50*); *σ*_21_ = 0.8 − 0.95 for cross immunity against mpox given smallpox infection (*51*) (*28*); ℛ_0,1_ = 1.5 − 2.7 for mpox; and *σ* = 12 = 0.1 − 0.9 for cross immunity against smallpox given mpox infection (which was assumed a wide range to explore large uncertainties).

##### S4.9 Within-subtype seasonal influenza model

Niche and fitness differences for within-subtype seasonal influenza strains are calculated by extrapolating cross immunity from antigenic distances among observed H3N2 influenza strains. First, we calculate the antigenic cartography based on 79 antisera and 273 viral isolates that were used in (*24*) using Racmacs package in R (*52*). Then, we take the positions of HK/68, EN/72, VI/75, TX/77, BK/79, SI/87, BE/89, BE/92, WU/95, SY/97, and FU02 strains from the resulting map and calculate their antigenic distances—for simplicity, we only consider the distances among consecutive pairs. Then, following (*27*) and (*53*), we model risk of reinfection 1−σ by multiplying antigenic distance *d* with a scaling factor *s* = 0.07, where *σ* represents cross immunity. Then, assuming that all strains have the same ℛ_0_ = 1.8, the corresponding mean susceptibility at endemic equilibrium can be calculated analytically:

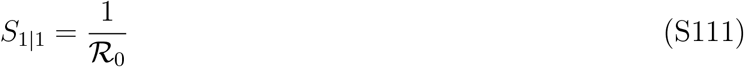

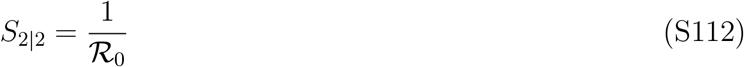

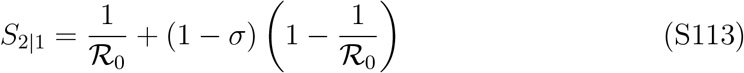

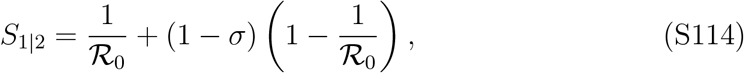

which allows us to compute the niche and fitness differences among different influenza strains. We note that most, major influenza evolution models assume (1) influenza strains have same ℛ_0_ and (2) infection with each strain provides life-long immunity (*31, 27, 53*). Therefore, niche and fitness differences can be calculated analytically for these models, based on the above equations.

##### S4.10 SARS-CoV-2 model

The dynamics of SARS-CoV-2 variants are modeled using our discrete-time adaptation of the predictive fitness model that was used by (*15*). The model has been previously validated against multinational data on SARS-CoV-2 variant dynamics (*15*). Briefly, the model seeks to capture the effects of immunity, either caused by natural infections or vaccination, on fitness of each variant, where the fitness is measured by the exponential growth rate. The model assumes that effective reproduction number of a given variant *i* exponentially decreases with immunity effect *C*_*ij*_(*t*):

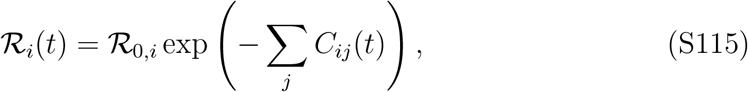

where *C*_*ij*_(*t*) represents the effect of infection or vaccination *j* on variant fitness at time *t*. This fitness effect is modeled as a function *H*(*T*_*ij*_) of a neutralization titer *T* :

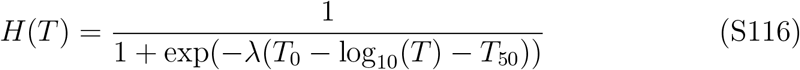

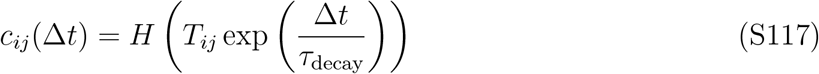

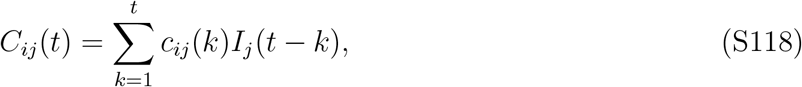

where *λ* = 3, *T*_0_ = log_10_(94), and *T*_50_ = log_10_(0.2 *×* 94) represent constant values for scaling the observed titers; *τ*_decay_ = 90 represents the scaling factor for exponential decay rate of antibody titers; and *c*_*ij*_(Δ*t*) represents the cross immunity effect of variant/vaccination *j* against variant *i* for Δ*t* days since infection/vaccination.

Then, assuming a fixed generation-interval distribution, variant fitness (i.e., epidemic growth rate) corresponds to:

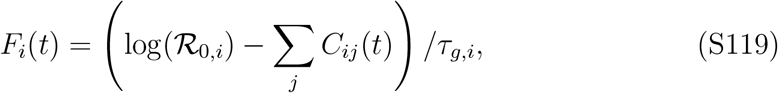

where *τ*_*g,i*_ represents the mean generation interval. The original model further incorporated a scaling factor that are multiplied with *C*_*ij*_(*t*) to account for under-reporting. Since we are simulating infections and all infections are known, we did not incorporate this scaling term. All parameters were taken from the corresponding GitHub repository (https://github.com/m-meijers/Population_Immunity_SARSCOV2) of the original article (*15*) due to differences in model parameterizations.

Since calculating niche and fitness difference only requires equilibrium conditions of one variant at a time, we rely on a single-strain model for these calculations. In the absence of vaccination, variant fitness *F* (*t*) and incidence of infection at time *I*(*t*) is given by

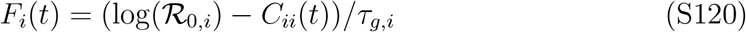

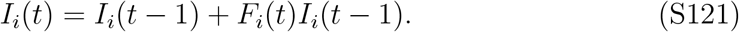

Then, the mean susceptibility at equilibrium is calculated as follows:

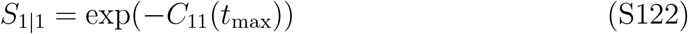

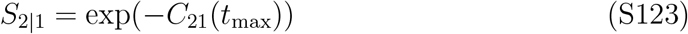

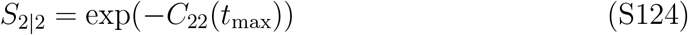

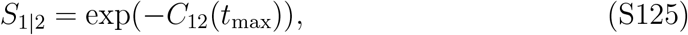

where *t*_max_ is set to 50 years.

To account for the effect of vaccination, we simply include vaccination terms in calculating the variant fitness:

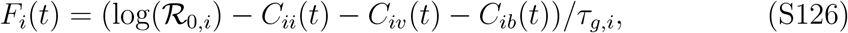

where *C*_*iv*_(*t*) and *C*_*ib*_(*t*) represent the effect of primary vaccination and booster series on fitness of variant *i*. To emulate a scenario in which a certain fraction of the population is constantly vaccinated, we assume

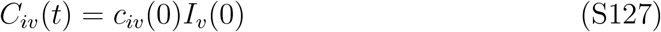

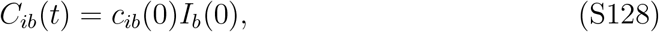

where *I*_*v*_(0) and *I*_*b*_(0) corresponds to vaccine coverage, and *c*_*iv*_(0) and *c*_*ib*_(0) represent the vaccinal cross immunity prior to any waning, which means that the estimated effects assume maximum amount of protection. Assuming more realistic changes in the incidence of vaccination will therefore result in weaker effects. Finally, the mean susceptibility at equilibrium is calculated as follows:

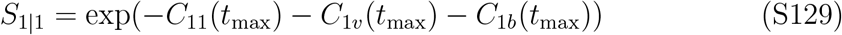

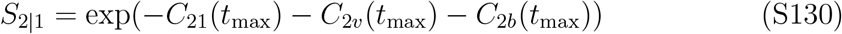

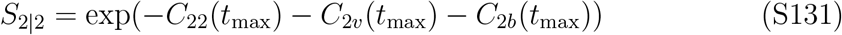

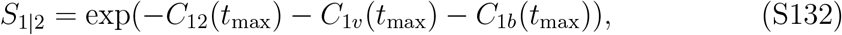

where *t*_max_ is set to 50 years.

For invasion simulations, we use a two strain model:

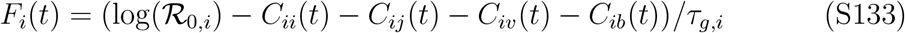

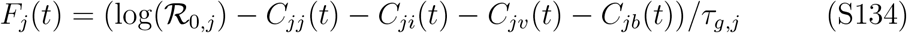

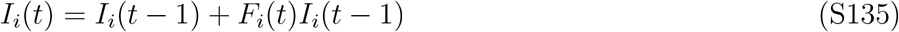

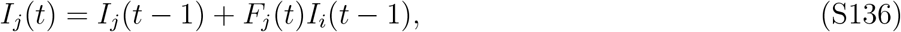

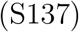

where *C*_*ij*_(*t*) and *C*_*ij*_(*t*) are defined as previously.

Based on (*15*), we assume the following for our base parameterization: *T*_11_ = 1, *T*_21_ = 1.8, *T*_22_ = 1, *T*_12_ = 1.8, *τ*_*g*,1_ = 7.8days, and *τ*_*g*,2_ = 5.5days for the WT and Alpha pair; *T*_11_ = 1, *T*_21_ = 2.8, *T*_22_ = 1, *T*_12_ = 3.8, *τ*_*g*,1_ = 5.5days, and *τ*_*g*,2_ = 3.5days for the Alpha and Delta pair; *T*_11_ = 1, *T*_21_ = 27, *T*_22_ = 1, *T*_12_ = 27, *τ*_*g*,1_ = 3.5days, and *τ*_*g*,2_ = 3days for the Delta and BA.1 pair; and *T*_11_ = 1, *T*_21_ = 2.4, *T*_22_ = 1, *T*_12_ = 3.6, *τ*_*g*,1_ = 3days, and *τ*_*g*,2_ = 3days for the BA.1 and BA.2 pair. The original code provides estimates of initial growth rate advantages *F*_*j*_(0) −F_i_(0): 0.052 for the WT and Alpha pair; 0.027 for the Alpha and Delta pair; 0.057 for the Delta and BA.1 pair; and 0.074 for the BA.1 and BA.2 pair. We translate this into ℛ_0_ estimates for each variant by assuming a fixed generation interval (to remain consistent with the original study) based on ℛ_0_ = 2.5 for the WT variant (*54*).

While the original model accurately captures the growth rate advantage of each variant but tends to give low estimates of the transmission advantage. Specifically, assuming ℛ_0_ = 2.5 for the wildtype variant, the original parameterization translates to ℛ_0_ = 3.33 for the Alpha variant, ℛ_0_ = 3.31 for the Delta variant, and ℛ_0_ = 3.77 for the BA.1 variant. We note that the Delta variant can still have a growth rate advantage compared to the Alpha variant due to its shorter generation interval despite its slightly lower ℛ_0_. On the other hand, other studies have suggested that the Delta variant is likely to be more transmissible than the Alpha variant across a wide range of assumptions about underlying generation-interval distributions (*55,34*). Therefore, we further explore scenarios assuming higher transmission advantages in Figure S2. Specifically, we assume 70%, 50%, 50% and 50% increases in ℛ_0_ for the WT vs Alpha pair (*37*), Alpha vs Delta pair (*55, 34*), Delta vs BA.1 pair (*33, 56, 57*), and BA.1 vs BA.2 pair, respectively (Figure S2).

The original parameterization further assumes that Omicron infections confer negligible (≈ 10%) protection against the Delta variant (and vice versa). This estimate is considerably lower than other estimates of cross protection (*33, 56, 57*).

Therefore, we performed additional sensitivity analysis by considering a wide range of values for cross protection (20%–60%) between Delta and BA.1 variants (assuming symmetric values of cross protection). This is done by setting *T*_*ij*_ to match *c*_*ij*_(0) to a given value of cross immunity.

#### S5 Effects of immunological details on the extent of overcompensation

In order to study how immunological details affect the extent of overcompensation and further shape competitive outcomes, we use a standard RSV model (*25*) and perform invasion simulations across a wide range of assumptions about transmission advantage of an invading variant and the duration of immunity. Specifically, we vary the mean duration of immunity between 1 to 20 years and the ratio of the ℛ_0_ of the invading variant and ℛ_0_ of the resident variant between 1 and 1.8. All other parameters are fixed to previously estimated values for RSV (*25*), and seasonality is not accounted for to simplify calculations. For each parameter combination, we compute niche and fitness differences to evaluate mutual invasibility and further perform an invasion simulation, which assumes that the resident variant is at equilibrium. We run the model for 10 years after the invasion of a new variant and evaluate the persistence of both resident and invading variants based on a prevalence cutoff of 10^−7^.

For each simulation, we also compute the effective reproduction number of the resident variant (i.e., variant 2, in this case):

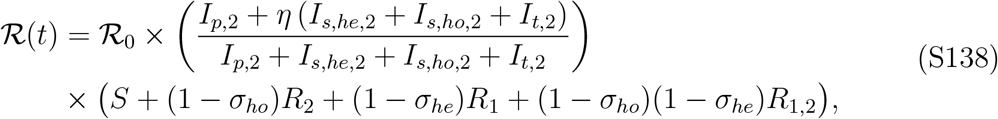

where ℛ_0_ represents the basic reproduction number; the second term captures reduction in transmissibility; and the last term captures reduction in susceptibility. Based on the effective reproduction number, we estimate the degree of overcompensation and return time. The degree of overcompensation is measured by 1 − min(ℛ(*t*)), and return time is measured by the time between the first time ℛ_eff_ decreases below 0.99 and the first time ℛ(*t*) increases above 0.99 again.

#### S6 Pneumococcal serotype simulations

We developed a deterministic approximation of the agent-based model used by (*11*) to estimate niche and fitness differences among pneumococcal serotypes. The original model considered three different mechanisms of serotype interactions: (1) immediate exclusion by the most fit resident, which reduces susceptibility against infections with other serotypes when the host is currently infected with another serotype; (2) serotype-specific immunity, which reduces susceptibility against reinfections with the same serotype after recovery, and (3) non-specific immunity, which reduces the duration of infection based on the total number of past colonization events, independent of serotypes.

Since calculating niche and fitness difference only requires equilibrium conditions of one variant at a time, we rely on a single-strain model for these calculations. The deterministic, single-strain dynamics of serotype *i* is then described by the following set of equations:

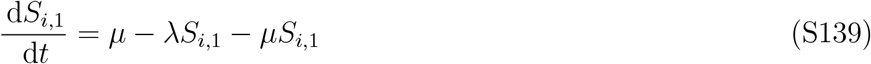

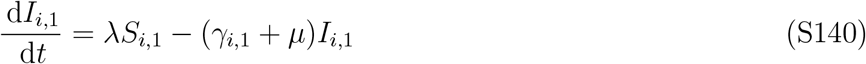

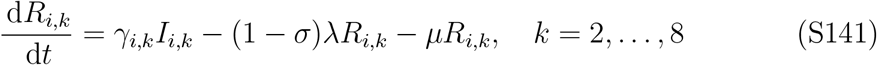

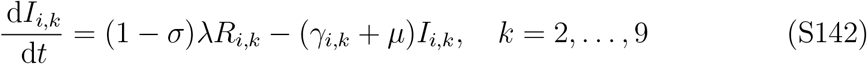

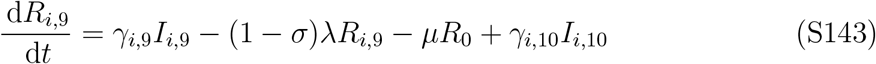

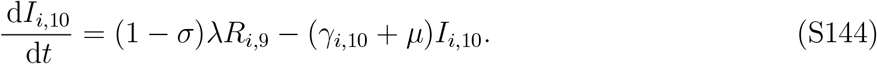

Here, *S*_*i*,1_ represents the proportion of immunologically naive individuals, *I*_*i,k*_ represents the proportion of infectious individuals who has previously experienced *k* − 1 infections; and *R*_*i,k*_ represents the proportion of partially immune individuals who has previously experienced *k* infections. The original model kept track of all infection history; for simplicity, we only count up to 9 infections and assume that any subsequent infections does not confer additional effect. Here, *λ* = *β*( ∑_*k*_ *I*_*i,k*_) represents the force of infection; a*σ* represents serotype-specific immunity. and *γ*_*i,k*_ represents the clearance rate, which is modeled as a function of number of past infections *k* − 1:

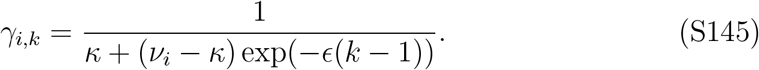

Here, *ν*_*i*_ represents the maximum duration of infection for serotype *i, κ* represents the minimum duration of infection, and ϵ represents the strength of non-specific immunity. Finally, *µ* represents the birth and death rates.

Then, mean intraspecific susceptibility at equilibrium is calculated as follows:

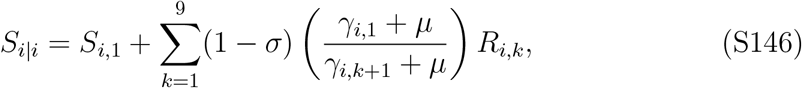

where (*γ*_*i*,1_ +*µ*)*/*(*γ*_*i,k*+1_+*µ*) captures reduction in duration of infection caused by non-specific immunity (see Section S3). Likewise, the mean interspecific susceptibility at equilibrium is calculated as follows:

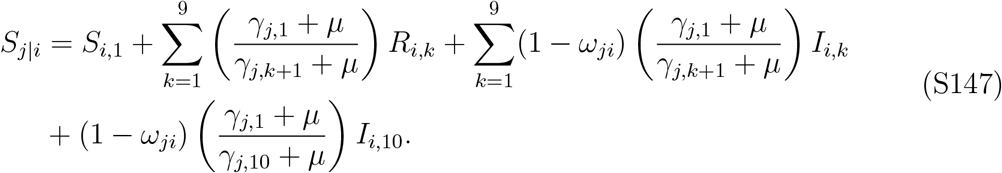

There are several key distinctions between intraspecific and interspecific susceptibility. First of all, serotype-specific immunity *σ* only affects the susceptibility against the same serotype and therefore does not affect susceptibility against different serotypes. Second, non-specific immunity affects all infections, and therefore reduces the duration of infection with serotype *j* depending on the number of past infections with serotype *i*—this is captured by the term (*γ*_*j*,1_+*µ*)*/*(*γ*_*j,k*+1_+*µ*). Finally, *ω*_*ji*_ = *µ*_max_(1 − (rank(i) − 1)/(Z − 1)) captures the effect of current infection with serotype *i* on susceptibility against infection with serotype *j*, where *µ*_max_ represents the maximum exclusion effect, rank(*i*) represents the fitness ranking of serotype *i* (with 1 being the most fit), and *Z* is the total number of serotypes being considered.

Following (*11*), we consider competition between *Z* = 25 serotypes, where we quantify niche and fitness different among all 300 pairs. We assume that the duration of infection linearly increases from 25 days to 220 days across 25 serotypes and set *β* = 1.1*/*(25*/*7)*/*weeks such that the least fit serotype has ℛ_0_ = 1.1. Other parameter values are given as follows: *κ* = 25*/*7 weeks, *ϵ* = 0.4, *σ* = 0.3, *µ* = 1*/*80*/*52*/*weeks, and *µ*_max_ = 0.25.

#### S7 Seasonal influenza simulations

We developed a semi-stochastic multistrain model by extending (*30*). The original work assumed a continuous evolution across a one-dimensional antigenic space, where the strength of cross immunity depends on antigenic distances between two strains.

Instead, we introduce stochastic jumps across the antigenic space, through which a new strain is introduced at the beginning of every season. Specifically, in the beginning of every season, the antigenic distance between the new and previous strains is randomly drawn from a gamma distribution with a mean of 1.3 units and a standard deviation of 0.2 units. Then, the underlying dynamics of each strain are solved deterministically every year based on (*30*):

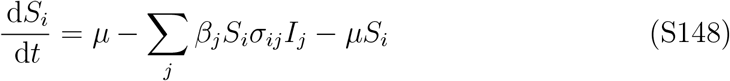

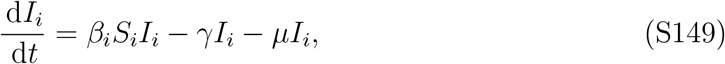

where *S*_*i*_ and *I*_*i*_ represent the proportion of individuals who are susceptible to and infected with strain *i*, respectively. Unlike standard compartmental models described previously, which keep track of infection history, this model keeps track of immune statuses, assuming that infection leads to either complete or no protection against future infection.

## Additional results

### S8 Impact of vaccination on mutual invasibility of SARS-CoV-2 variants

Vaccination represents an important perturbation of pathogen fitness and has had significant effect in shaping evolutionary trajectories of SARS-CoV-2 variants (*15*). When viewed through the lens of PIT, an increase in overall population-level immunity corresponds to a decrease in niche difference (Figure S1), but asymmetric effects of vaccination against different variants (e.g., generally providing stronger protection against variants that are more similar to the WT variant) can result in an increase in fitness difference. However, these effects are not strong enough to rule out mutual invasion, except for the WT, Alpha pair. Even when we assume greater transmission advantages between successive variants, we still find that all variant pairs, except for the WT and Alpha, can mutually invade one another (Figure S2). A more detailed sensitivity analysis for the Delta, BA. 1 pair shows that mutual invasion is universal across a wide range of assumptions (Figure S3). We find that preventing mutual invasion requires: (1) strong cross immunity (*>* 60%), (2) high transmission advantage of the BA. 1 variant (*×* 2–3), (3) basic reproduction number around 3–4 for the Delta variant, and (4) sufficient levels of booster vaccination. In all other cases, mutual invasion is still allowed.

### S9 Case study: quantifying coexistence mechanisms for pneu-moccocal serotypes

To illustrate this utility of our framework, we re-visit coexistence patterns of pneumococcal serotypes, a core example of examining co-circulation in infectious disease pathogens (*11*). Specifically, (*11*) previously showed that the diversity of pneumococcal serotypes, despite large differences in their duration of infection, are maintained through a combination of serotype-specific immunity and non-specific immunity. They further argued that serotype-specific immunity is a stabilizing mechanism, which increases niche difference (and therefore reduces niche overlap) and non-specific immunity is an equalizing mechanism, which decreases the fitness difference. The SIS-like nature of pneumococcal serotypes, which permits multiple homologous reinfections, minimizes the degree of overcompensation, making it an ideal candidate for applying our framework to tease apart coexistence mechanisms; for this system, pair-wise predictions of coexistence can be also reliably extended to predictions about coexistence in a multistrain system.

Our analysis offers a more detailed explanation (Figure S4). In fact, serotype-specific immunity alone has a negligible effect on both niche (Figure S4A) and fitness difference (Figure S4B). On the other hand, non-specific immunity reduces the duration of infection for all serotypes depending on the number of past infections from any serotype, which not only decreases the fitness difference (Figure S4B; because it limits their transmission) but also decreases niche difference (Figure S4A; because previously infected individuals are effectively protected from future infections). When serotype-specific immunity is added in the presence of non-specific immunity, serotype-specific immunity reduces the amount of reinfection, which in turn limits the overall effect of non-specific immunity, thereby recovering some of the fitness difference (Figure S4B) as well as the niche difference (Figure S4A). These changes are further plotted in Figure S4C, where each point indicates a pairwise interaction between two serotypes among any two 25 serotypes. For example, serotype 1, which has a higher reproduction number, competitively excludes serotype 25 in the baseline scenario, but they can mutually invade each other and co-circulate in the presence of both serotype-specific and non-specific immunity. Overall, our analysis highlights that the relative strengths of serotype-specific and non-specific immunity determines the competitive outcomes of pneumoccocal serotypes.

### S10 Invasion simulations for SARS-CoV-2 variants

Performing invasion simulations for successive variant pairs correctly predicts that the invasion of a novel variant results in an extinction of resident strain for the WT, Alpha pair as well as the BA.1 and BA.2 pairs (Figure S5). However, the simulations predict that the invasion of Delta variant could lead to a stable co-circulation whereas the invasion of the Omicron variant could lead to an extinction of the BA.1 variant and persistence of the Delta variant (Figure S5); these results do not match the observed patterns of strain replacement.

The latter predictions depends on the original parameterization by (*15*). As explained earlier, the original parameterization tends to underestimate the transmission advantage. When we conservatively assume that the Delta variant is 50% more transmissible than the Alpha variant (assuming ℛ_0_ = 4.25 for the Alpha variant based on 70% increased transmissibility (*37*)), we find that the invasion of the Delta variant can lead to extinction of both Alpha and Delta variants with or without vaccination (Figure S6). We note that Alpha and Delta variants can still mutually invade under this assumption, as shown in Figure S2.

Further assuming a 50% increased transmissibility of the BA.1 variant (i.e., 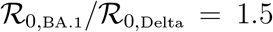 still leads to persistence of the Delta variant following the invasion of the Omicron variant (Figure S7). This is also due to the original parameterization by (*15*), which assumes that Omicron infections confer negligible (≈ 10%) protection against the Delta variant (and vice versa). When we assume a 40% cross protection (*33, 56, 57*) and a 50% increased transmissibility of the BA.1 variant, we find that the invasion of the BA.1 variant can lead to an extinction of both Delta (resident) and BA.1 (invading) variants (Figure S7).

We note that there are many other complexities that likely contributed to the strain replacement of SARS-CoV-2 variants. Most importantly, none of the variants were near equilibrium, and so their prevalence during the invasion of a novel variant was likely considerably lower than the equilibrium prevalence, which would have made extinction through overcompensation easier. As illustrated in influenza simulations, history of past infections and vaccination likely contributed to strain replacement. Finally, the onset of non-pharmaceutical interventions in response to emergence of novel variants also likely contributed to extinction of resident variants.

### S11 Invasion simulations for seasonal influenza strains

To better understand the apparent lack of co-circulation among strictly antigenically distant strains in a seasonal influenza subtype, we extended a previously developed multistrain model for influenza (*30*) by including stochastic strain jumps, which allowed us to capture key features of antigenic evolution of seasonal influenza strains (Figure S8A). As an example, for strains *x* and *y*, which correspond to the dominant strains in years 13 and 15 respectively, we find that both strains can mutually invade one another (Figure S8B,C). However, the immune escape of an invading strain permits a large outbreak, which in turn causes a large susceptible depletion (Figure S8D–F) and drives the resident strain to very low prevalence (*<* 10^−8^). In the presence of demographic stochasticity in a finite population, the resident strain would go extinct, and therefore, two strains will not be able to co-circulate.

In reality, invasion at endemic equilibrium rarely happens since most influenza strains become extinct in a few years. To study how immune profiles and the order of strain introduction affect coexistence patterns, we take the immune statuses at year 12.5 and sequentially introduce strains *x* and *y* on years 12.5 and 13.5 (Figure S9.

When strain *x* is introduced first, followed by strain *y*, both strains go extinct. This is because the population-level susceptibility is farther away from the equilibrium, which causes a large susceptible depletion. When strain *y* is introduced first, it causes an even greater susceptible depletion, which then leads to an extinction of strain *y* and also prevents strain *x* from invading.

We further evaluate the impact of overcompensatory dynamics on co-circulation patterns by running invasion simulations across a wide range of assumptions about cross immunity and pathogen transmissibility using the same influenza model (Figure S8G). Although a wide array of parameters permit mutual invasion, the region of co-circulation is extremely narrow: in most cases, the invasion of a novel variant leads to extinction of both variants. We note that the multistrain model proposed by (*30*) implicitly assumes that individuals who have been infected with serotype *i* but remain susceptible to serotype *j* can still be exposed again by serotype *i* and, with some probability, become immune to serotype *j*. This implicit assumption causes an underestimation of mutual invasibility. Specifically, interspecific susceptibility at equilibrium for this model corresponds to

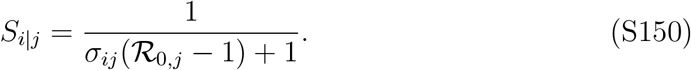

In contrast, for standard SIR models, interspecific susceptibility at equilibrium for this model corresponds to

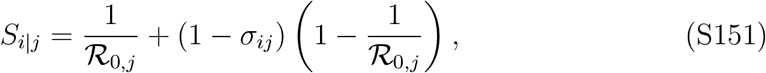

which provides an even wider region for mutual invasion. This example further provides support for near-universality of mutual invasibility.

Interestingly, introducing the invader near equilibrium susceptibility makes co-circulation much more likely, suggesting that repeated invasion of a novel variant (through mutation or spillover events) may promote co-circulation (Figure S10). Similar ideas have been discussed previously in the context of influenza evolution simulations to prevent influenza strains from going extinct during the initial introduction to the population (*27, 53*).

## Supplementary Figure

**Figure S1:**
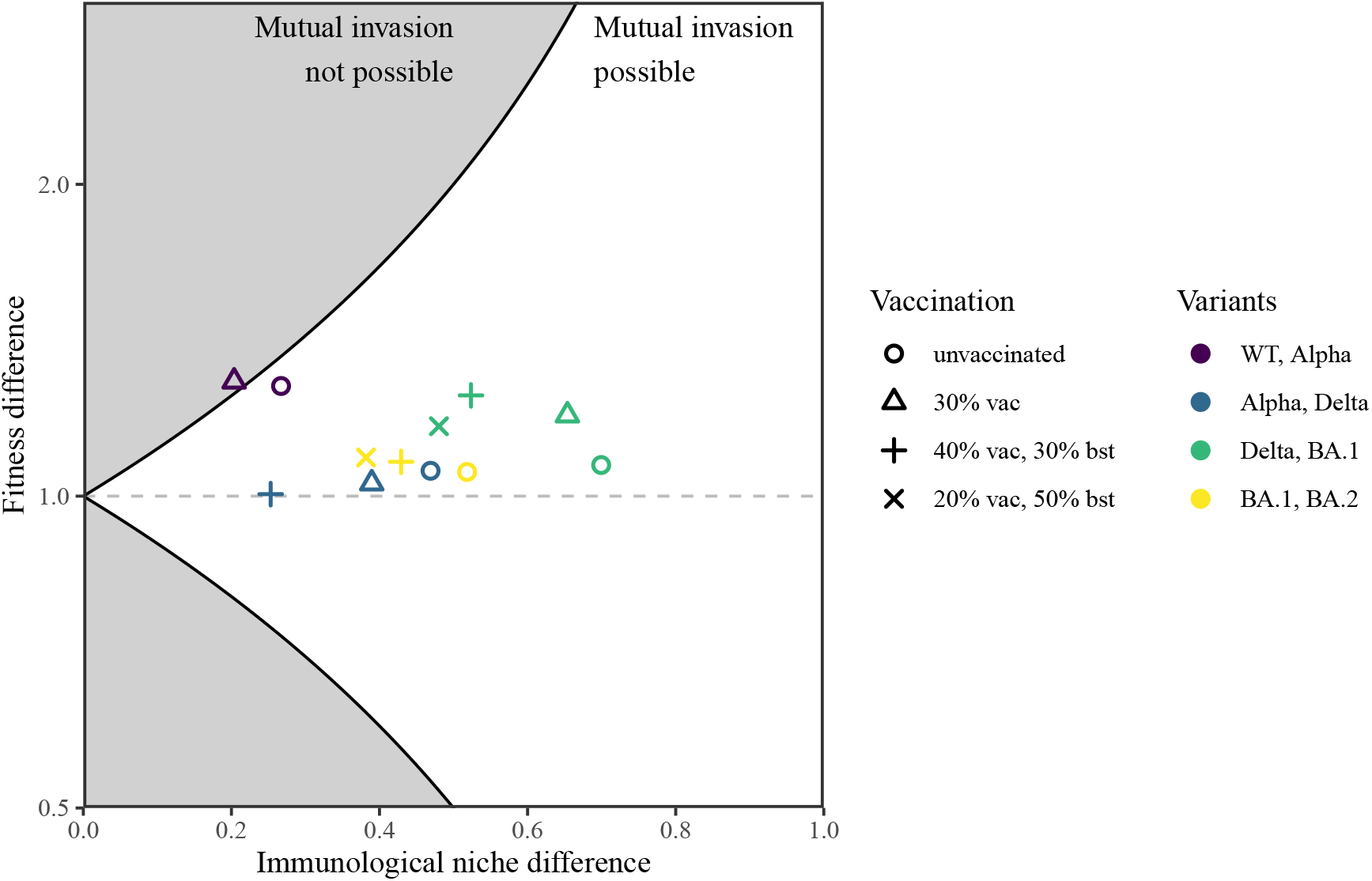
Impact of vaccination on mutual invasibility of SARS-CoV-2 variants. Niche and fitness difference estimates across successive SARS-CoV-2 variants under different vaccination scenarios with an increasing vaccine pressure. Points in the white region allow mutual invasion, meaning that a variant can invade a population when the competing variant is at equilibrium. Points in the grey region do not allow mutual invasion, meaning that the presence of more fit variant prevents the invasion of less fit variant. Two vaccination types are considered: primary series (“vac”) and booster shots (“bst”). For example, “40% vac, 30% bst” scenario represents a case where 40% of the total population has only received primary series and an additional 30% of the total population has received booster shots (therefore, a total coverage of 70%). Vaccine effectiveness and cross immunity values among SARS-CoV-2 variants are determined calculated from neutralization titers following (*15*). We plot max(*f*_*i*_*/f*_*j*_, *f*_*j*_*/f*_*i*_) such that fitness difference is always greater than 1. The fitness difference is plotted on a log scale for symmetry.

**Figure S2:**
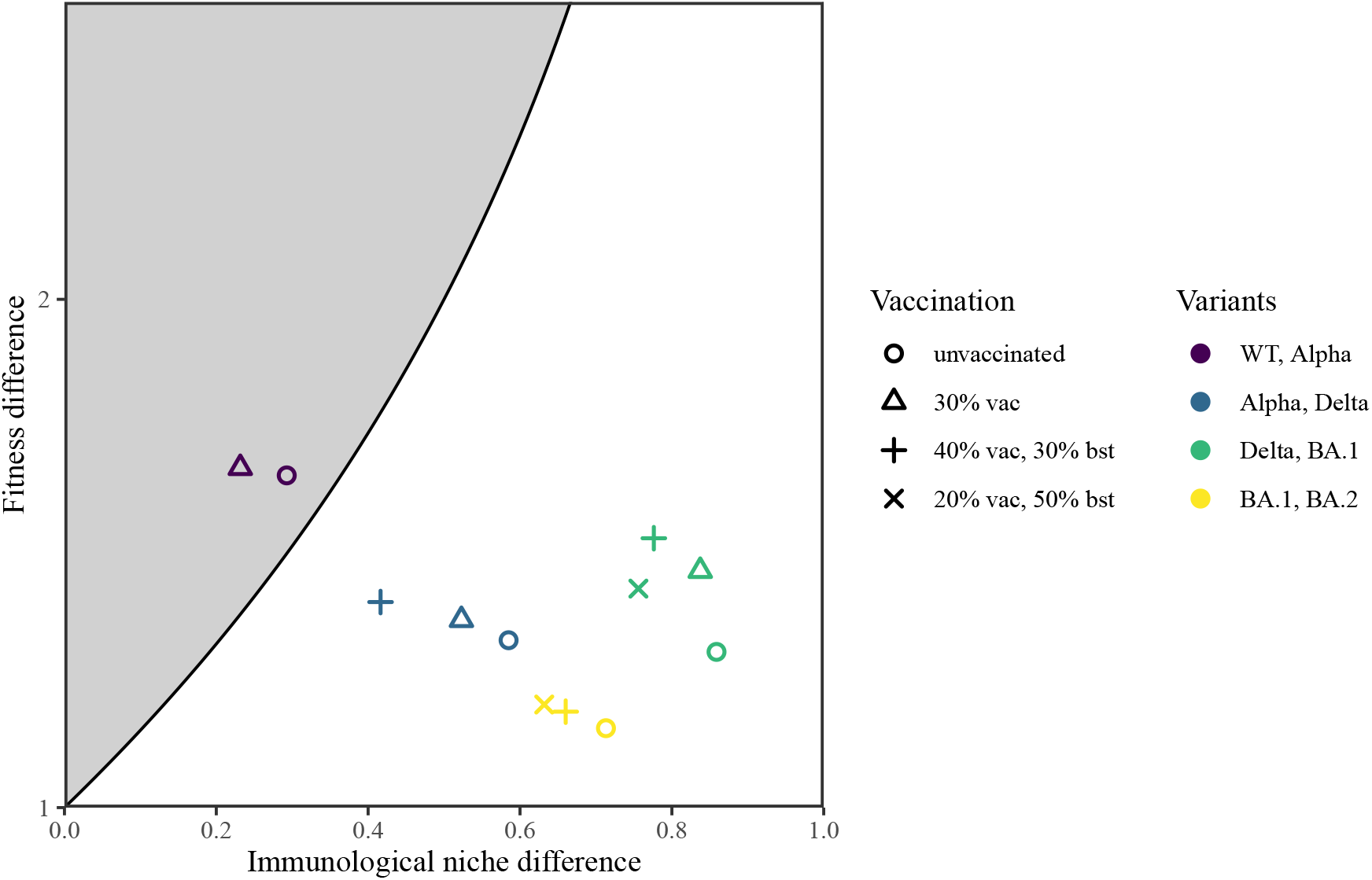
Sensitivity analysis for SARS-CoV-2 variants assuming higher transmission advantage. As explained earlier, the original parameterization by (*15*) tends to provide lower estimates of variant transmission advantage compared to other studies. Here, we assume 70%, 50%, 50% and 50% increases in *ℛ*_0_ for the WT vs Alpha pair (*37*), Alpha vs Delta pair (*55, 34*), Delta vs BA.1 pair (*33, 56, 57*), and BA.1 vs BA.2 pair, respectively. See Figure S1 for detailed captions.

**Figure S3:**
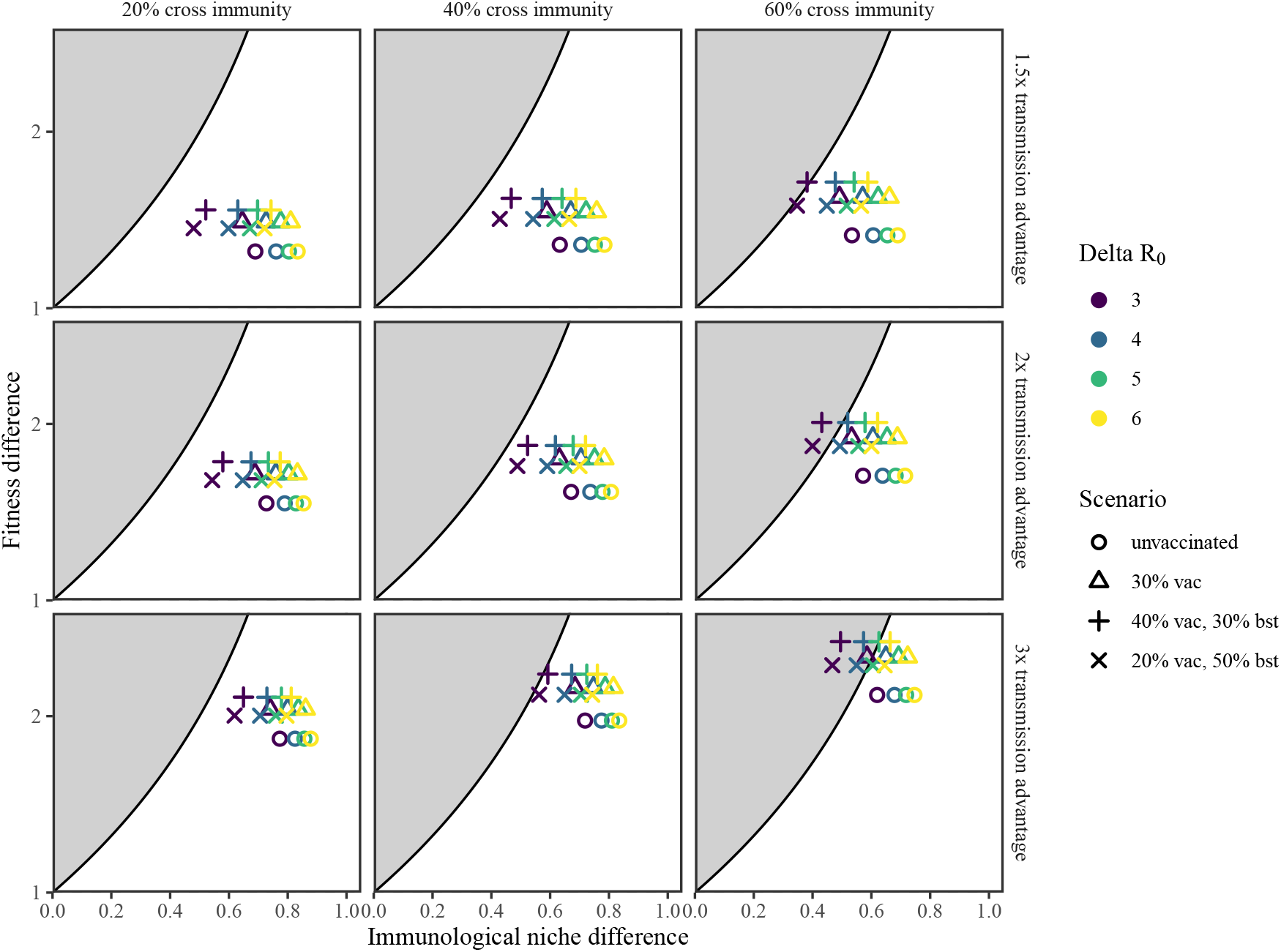
Sensitivity analysis for SARS-CoV-2 Delta and BA.1 variants across a wide range of scenarios. See Figure S1 for detailed captions.

**Figure S4:**
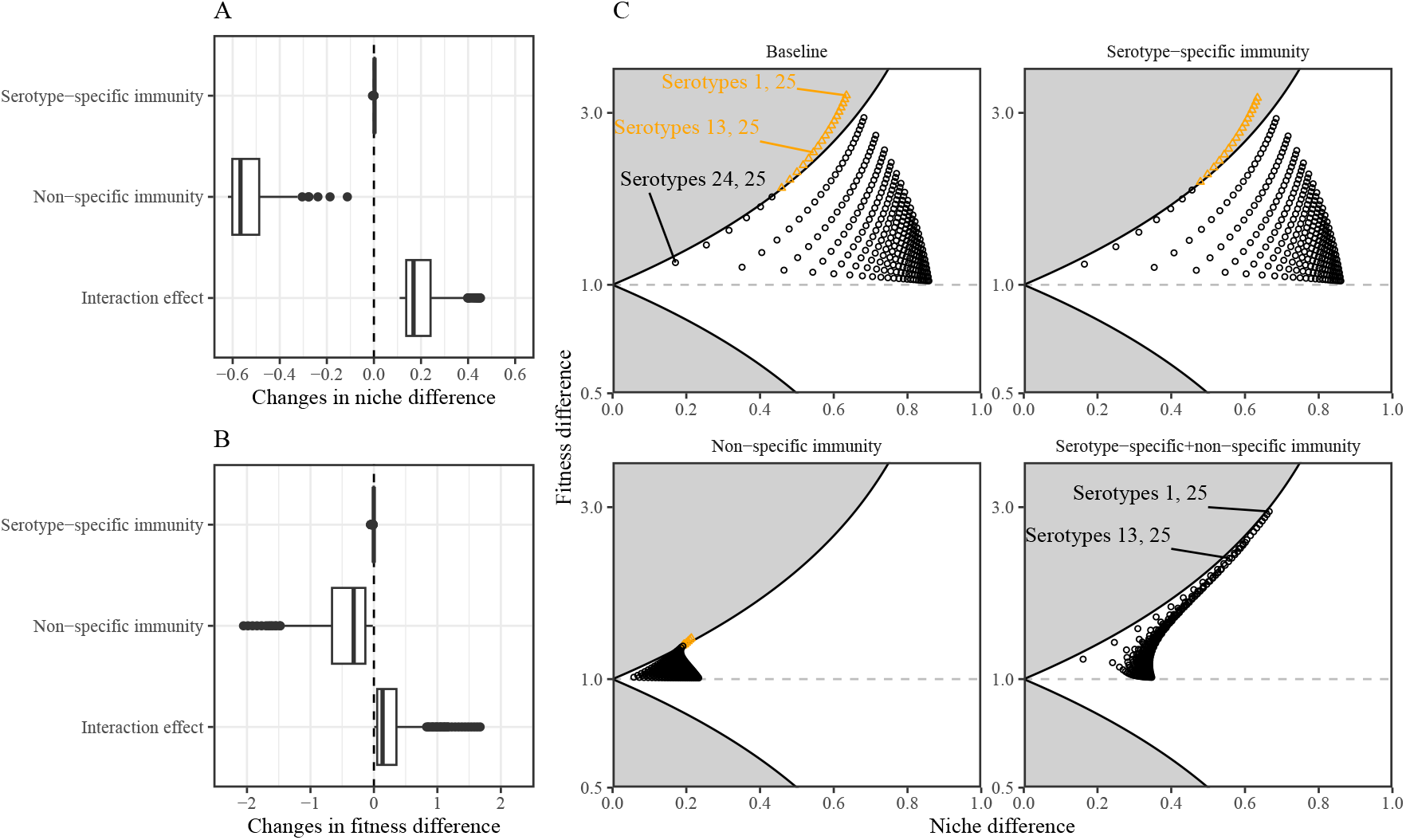
Quantifying coexistence mechanisms of pneumococcal serotypes. (A) Changes in niche overlap caused by serotype-specific immunity, non-specific immunity, and their interaction effects. (B) Changes in fitness difference caused by serotype-specific immunity, non-specific immunity, and their interaction effects. (C) Niche overlap and fitness difference estimates across all possible pairs among 25 pneumococcal serotypes under different immune mechanisms. Serotypes are numbered based on their reproduction (serotype 1 being the most transmissible and serotype 25 being the least transmissible). Black points (inside the white region) indicate serotype pairs that permit mutual invasion. Orange triangles (inside the gray region) indicate serotype pairs that result do not permit mutual invasion. Estimates of changes in niche overlap and fitness difference are calculated by comparing the resulting niche overlap and fitness difference estimates under each scenario with those under the baseline scenario. Interaction effects are calculated by subtracting two main effects from the total differences between “serotype-specific+non-specific immunity” scenario and the baseline scenario following the approach outlined in (*58*).

**Figure S5:**
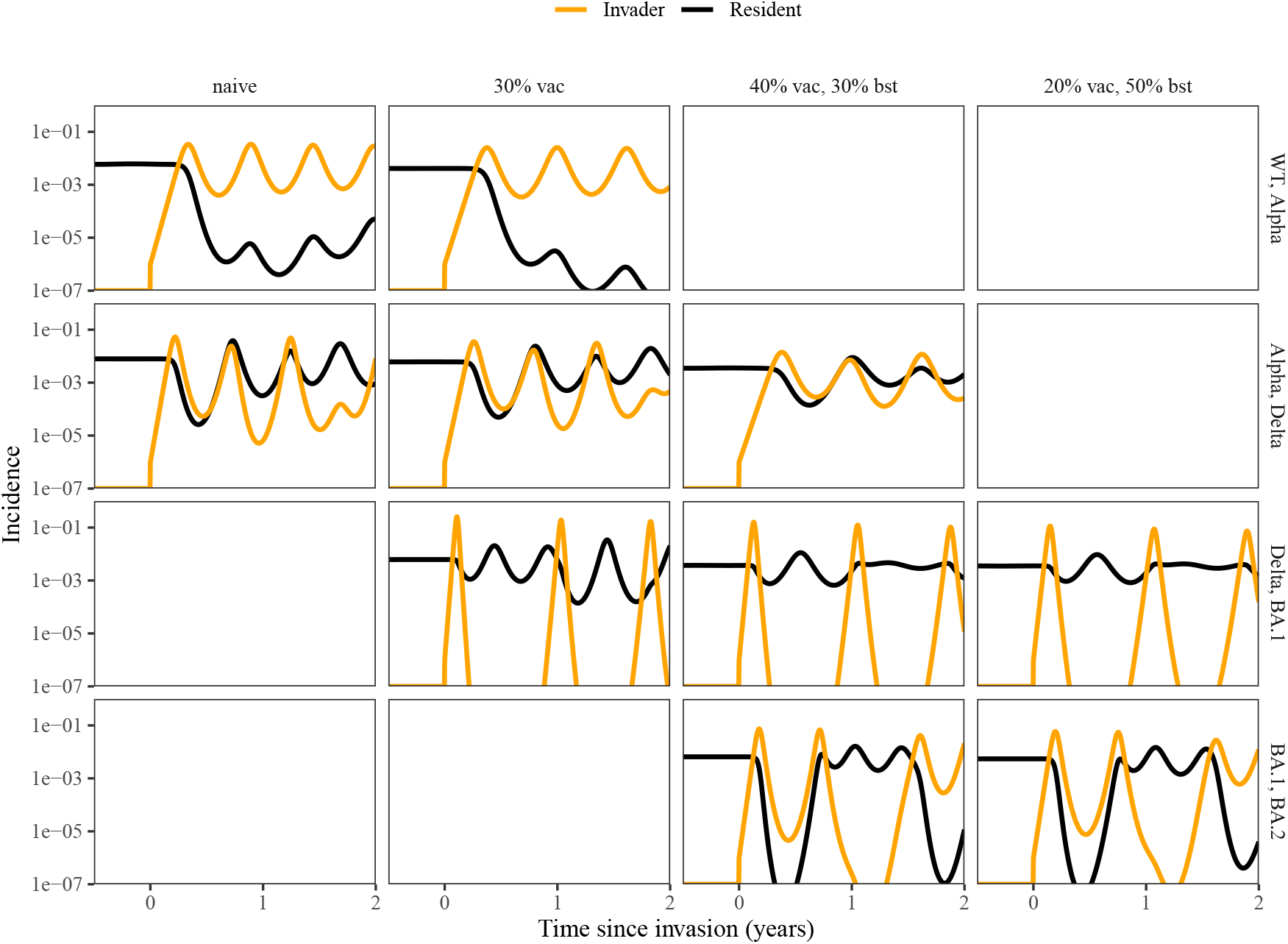
Invasion simulations for SARS-CoV-2 variants. Invasion simulations were performed based on the original parameterizations by (*15*). We only consider invasion simulations under vaccination scenarios that are relevant to the observed dynamics. Orange lines represent incidence trajectories for the invading variant. Black lines represent incidence trajectories for the resident variant. Two vaccination types are considered: primary series (“vac”) and booster shots (“bst”); see Figure S1 for detailed explanations on vaccination scenarios.

**Figure S6:**
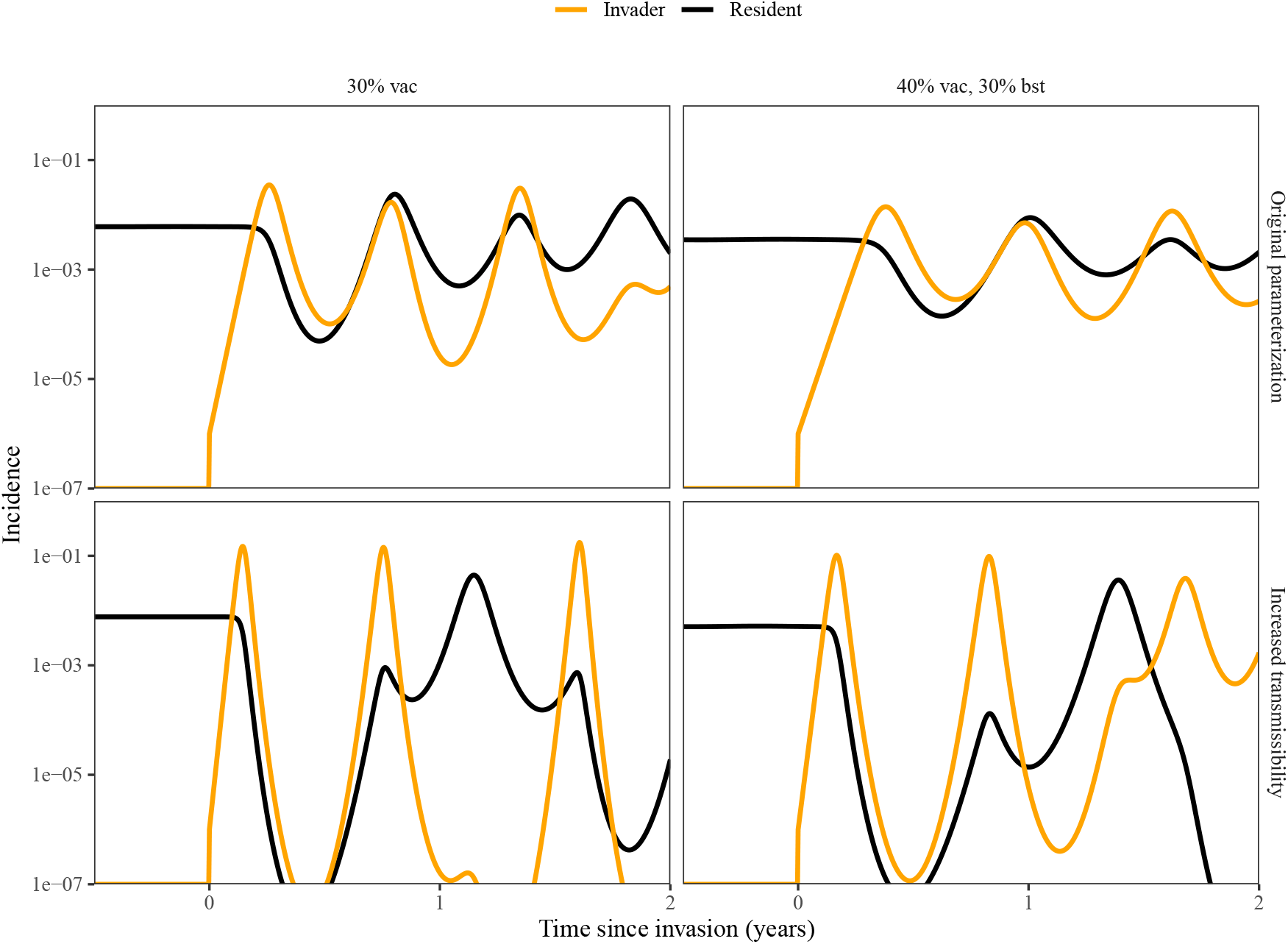
Invasion simulations for SARS-CoV-2 Alpha and Delta variants assuming increased transmissibility. Invasion simulations were performed assuming 50% increase in *ℛ*_0_ for the Delta variant compared to the Alpha variant (and further assuming 70% increase in *ℛ*_0_ for the Alpha variant compared to the WT variant). Orange lines represent incidence trajectories for the invading variant. Black lines represent incidence trajectories for the resident variant. Two vaccination types are considered: primary series (“vac”) and booster shots (“bst”); see Figure S1 for detailed explanations on vaccination scenarios.

**Figure S7:**
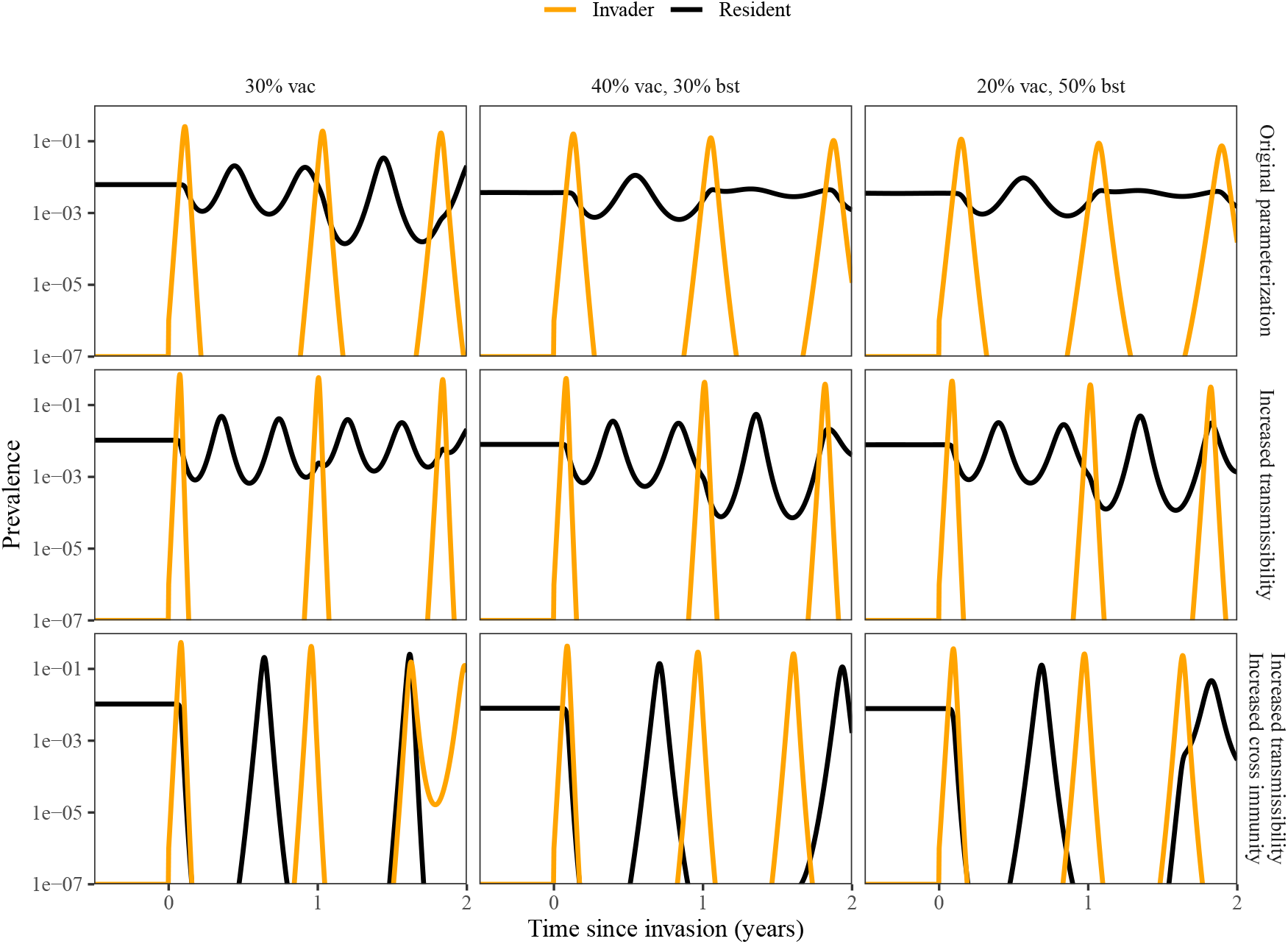
Invasion simulations for SARS-CoV-2 Delta and Omicron variants assuming increased transmissibility and cross immunity. Invasion simulations were performed assuming 50% increase in *R*_0_ for the BA.1 variant compared to the Delta variant (and further assuming 50% increase in *R*_0_ for the Delta variant and 70% increase for the Alpha variant). Orange lines represent incidence trajectories for the invading variant. Black lines represent incidence trajectories for the resident variant. Two vaccination types are considered: primary series (“vac”) and booster shots (“bst”); see Figure S1 for detailed explanations on vaccination scenarios.

**Figure S8:**
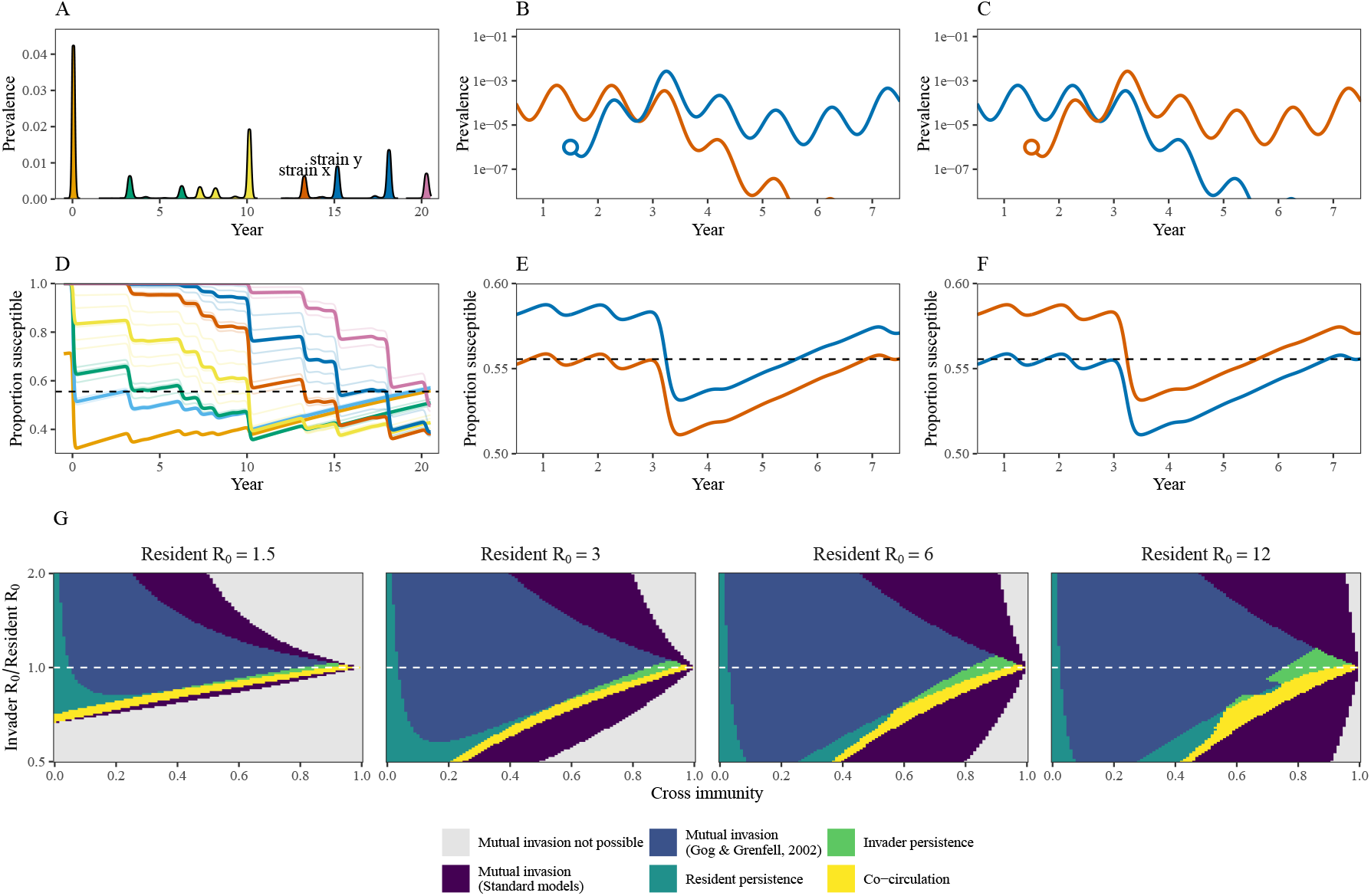
Influenza dynamics and mutual invasion simulations. (A) Time series of prevalence of influenza infections. Colors indicate different influenza subtypes. (B,C) Mutual invasion simulations for strains *x* (orange) and *y* (blue) when the resident strain is at equilibrium. (D–F) Corresponding susceptible dynamics to panels A–E. (G) 5 year co-circulation/persistence when resident is at equilibrium and invader is at rare. Mutual invasion not possible: The presence of a variant with higher ℛ_0_ prevents the invasion of a variant with lower ℛ_0_. Mutual invasion: both variants can mutually invade each another but the invasion of the invader leads to extinction of both variants. Resident persistence: invader becomes extinct after invasion, and the resident persists. Invader persistence: invader persists after invasion, but the resident becomes extinct. Co-circulation: both invader and resident persist after invasion. Persistence is measured by a prevalence cutoff of 10^−7^.

**Figure S9:**
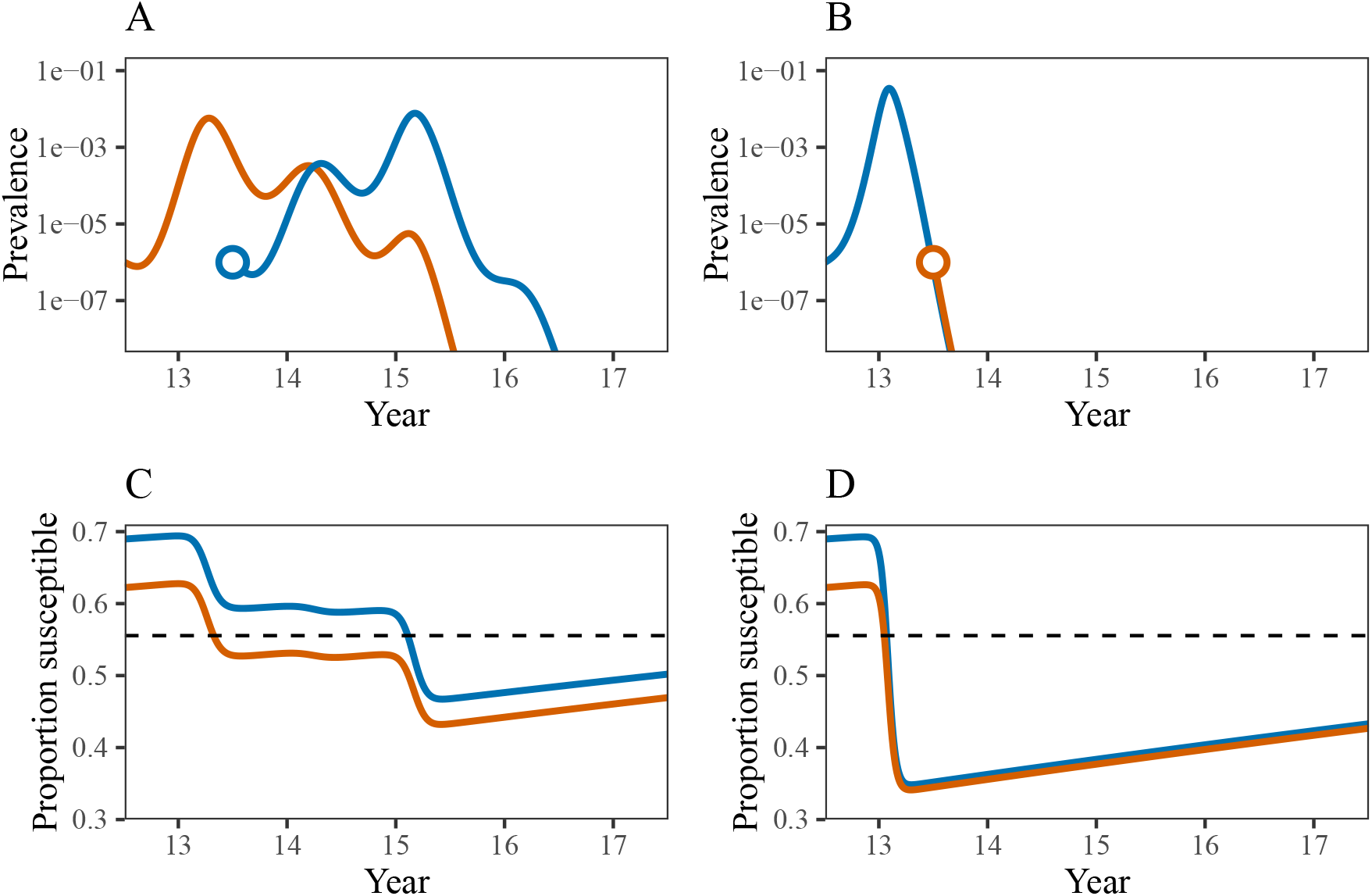
Invasion simulation when for strains *x* and *y* given immunity profile at year 12. (A,B) Mutual invasion simulations for strains *x* (orange) and *y* (blue). (C,D) Corresponding susceptible dynamics to panels A and B.

**Figure S10:**
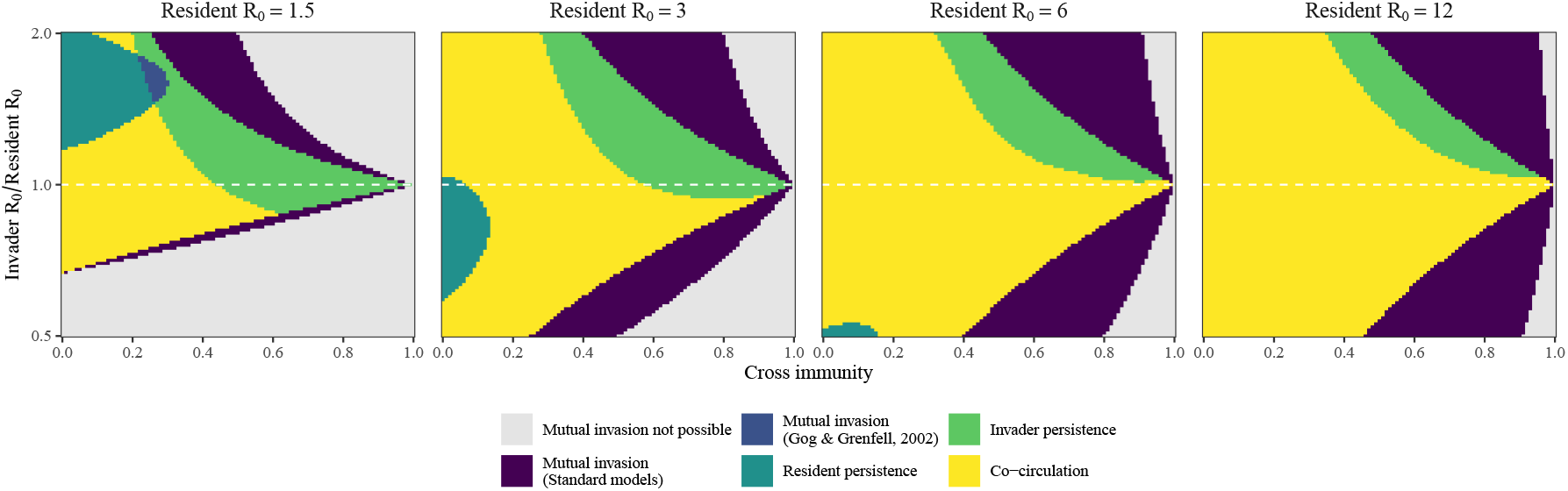
5 year coexistence/persistence when resident is at equilibrium and invader is near equilibrium. Effective reproduction number of the invading variant was set to 1.05 by adjusting the proportion of individuals who are susceptible to the invading variant. Mutual invasion not possible: The presence of a variant with higher ℛ_0_ prevents the invasion of a variant with lower ℛ_0_. Mutual invasion: both variants can mutually invade each another but the invasion of the invader leads to extinction of both variants. Resident persistence: invader becomes extinct after invasion, and the resident persists. Invader persistence: invader persists after invasion, but the resident becomes extinct. Co-circulation: both invader and resident persist after invasion. Persistence is measured by a prevalence cutoff of 10^−7^.

## Notes

https://github.com/parksw3/invasion

### Competing Interest Statement

The authors have declared no competing interest.

## References

1. P. Rohani, D. J. Earn, B. Finkenstädt, and B. T. Grenfell. Population dynamic interference among childhood diseases. Proc. R. Soc. B, 265(1410):2033–2041, 1998.

2. J. O. Lloyd-Smith. Vacated niches, competitive release and the community ecology of pathogen eradication. Philos. Trans. R. Soc. B, 368(1623):20120150, 2013.

3. E. W. Seabloom, E. T. Borer, K. Gross, A. E. Kendig, C. Lacroix, C. E. Mitchell, E. A. Mordecai, and A. G. Power. The community ecology of pathogens: coinfection, coexistence and community composition. Ecol. Lett., 18(4):401–415, 2015.

4. S. Nickbakhsh, C. Mair, L. Matthews, R. Reeve, P. C. D. Johnson, F. Thorburn, B. Von Wissmann, A. Reynolds, J. McMenamin, R. N. Gunson, and P. R. Murcia. Virus– virus interactions impact the population dynamics of influenza and the common cold. Proc. Natl. Acad. Sci. U.S.A., 116(52):27142–27150, 2019.

5. B. L. Rice, D. C. Douek, A. B. McDermott, B. T. Grenfell, and C. J. E. Metcalf. Why are there so few (or so many) circulating coronaviruses? Trends Immunol., 42(9):751–763, 2021.

6. A. J. Sieben, J. R. Mihaljevic, and L. G. Shoemaker. Quantifying mechanisms of coexistence in disease ecology. Ecology, 103(12):e3819, 2022.

7. M. Woolhouse and E. Gaunt. Ecological origins of novel human pathogens. Crit. Rev. Microbiol., 33(4):231–242, 2007.

8. N. M. Ferguson, A. P. Galvani, and R. M. Bush. Ecological and immunological determinants of influenza evolution. Nature, 422(6930):428–433, 2003.

9. O. Restif and B. T. Grenfell. Integrating life history and cross-immunity into the evolutionary dynamics of pathogens. Proc. R. Soc. B, 273(1585):409–416, 2006.

10. V. E. Pitzer, M. M. Patel, B. A. Lopman, C. Viboud, U. D. Parashar, and B. T. Grenfell. Modeling rotavirus strain dynamics in developed countries to understand the potential impact of vaccination on genotype distributions. Proc. Natl. Acad. Sci. U.S.A., 108(48):19353–19358, 2011.

11. S. Cobey and M. Lipsitch. Niche and neutral effects of acquired immunity permit coexistence of pneumococcal serotypes. Science, 335(6074):1376–1380, 2012.

12. N. G. Reich, S. Shrestha, A. A. King, P. Rohani, J. Lessler, S. Kalayanarooj, I.-K. Yoon, R. V. Gibbons, D. S. Burke, and D. A. T. Cummings. Interactions between serotypes of dengue highlight epidemiological impact of cross-immunity. J. R. Soc. Interface, 10(86):20130414, 2013.

13. S. Bhattacharyya, P. H. Gesteland, K. Korgenski, O. N. Bjørnstad, and F. R. Adler. Cross-immunity between strains explains the dynamical pattern of paramyxoviruses. Proc. Natl. Acad. Sci. U.S.A., 112(43):13396–13400, 2015.

14. S. M. Kissler, C. Tedijanto, E. Goldstein, Y. H. Grad, and M. Lipsitch. Projecting the transmission dynamics of SARS-CoV-2 through the postpandemic period. Science, 368(6493):860–868, 2020.

15. M. Meijers, D. Ruchnewitz, J. Eberhardt, M. Łuksza, and M. Lässig. Population immunity predicts evolutionary trajectories of SARS-CoV-2. Cell, 186(23):5151–5164, 2023.

16. G.-Y. Kim, I. Rheem, Y. H. Joung, and J. K. Kim. Investigation of occurrence patterns of respiratory syncytial virus A and B in infected-patients from Cheonan, Korea. Respir. Res., 21:1–9, 2020.

17. S. Takahashi, Q. Liao, T. P. Van Boeckel, W. Xing, J. Sun, V. Y. Hsiao, C. J. E. Metcalf, Z. Chang, F. Liu, J. Zhang, J. T. Wu, B. J. Cowling, G. M. Leung, J. J. Farrar, H. R. van Doorn, B. T. Grenfell, and H. Yu. Hand, foot, and mouth disease in China: modelling epidemic dynamics of enterovirus serotypes and implications for vaccination. PLOS Med., 13(2):e1001958, 2016.

18. L. C. Katzelnick, L. Gresh, M. E. Halloran, J. C. Mercado, G. Kuan, A. Gordon, A. Balmaseda, and E. Harris. Antibody-dependent enhancement of severe dengue disease in humans. Science, 358(6365):929–932, 2017.

19. N. Ferguson, R. Anderson, and S. Gupta. The effect of antibody-dependent enhancement on the transmission dynamics and persistence of multiple-strain pathogens. Proc. Natl. Acad. Sci. U.S.A., 96(2):790–794, 1999.

20. D. Gökaydin, J. B. Oliveira-Martins, I. Gordo, and M. G. M. Gomes. The reinfection threshold regulates pathogen diversity: the case of influenza. J. R. Soc. Interface, 4(12):137–142, 2007.

21. D. Zinder, T. Bedford, S. Gupta, and M. Pascual. The roles of competition and mutation in shaping antigenic and genetic diversity in influenza. PLOS Pathog., 9(1):e1003104, 2013.

22. L. Yan, R. A. Neher, and B. I. Shraiman. Phylodynamic theory of persistence, extinction and speciation of rapidly adapting pathogens. eLife, 8:e44205, 2019.

23. P. Chesson. Mechanisms of maintenance of species diversity. Annu. Rev. Ecol. Evol. Syst., 31(1):343–366, 2000.

24. D. J. Smith, A. S. Lapedes, J. C. De Jong, T. M. Bestebroer, G. F. Rimmelzwaan, A. D. M. E. Osterhaus, and R. A. M. Fouchier. Mapping the antigenic and genetic evolution of influenza virus. Science, 305(5682):371–376, 2004.

25. L. J. White, M. Waris, P. A. Cane, D. J. Nokes, and G. F. Medley. The transmission dynamics of groups A and B human respiratory syncytial virus (hRSV) in England & Wales and Finland: seasonality and cross-protection. Epidemiol. Infect., 133(2):279–289, 2005.

26. K. Koelle, M. Pascual, and M. Yunus. Serotype cycles in cholera dynamics. Proc. R. Soc. B, 273(1603):2879–2886, 2006.

27. T. Bedford, A. Rambaut, and M. Pascual. Canalization of the evolutionary trajectory of the human influenza virus. BMC Biol., 10:1–12, 2012.

28. R. Grant, L.-B. L. Nguyen, and R. Breban. Modelling human-to-human transmission of monkeypox. Bull. World Health Organ., 98(9):638, 2020.

29. W. Yang, E. H. Y. Lau, and B. J. Cowling. Dynamic interactions of influenza viruses in Hong Kong during 1998–2018. PLOS Comput. Biol., 16(6):e1007989, 2020.

30. J. R. Gog and B. T. Grenfell. Dynamics and selection of many-strain pathogens. Proc. Natl. Acad. Sci. U.S.A., 99(26):17209–17214, 2002.

31. K. Koelle, S. Cobey, B. Grenfell, and M. Pascual. Epochal evolution shapes the phylodynamics of interpandemic influenza A (H3N2) in humans. Science, 314(5807):1898–1903, 2006.

32. T. Bedford, M. A. Suchard, P. Lemey, G. Dudas, V. Gregory, A. J. Hay, J. W. McCauley, C. A. Russell, D. J. Smith, and A. Rambaut. Integrating influenza antigenic dynamics with molecular evolution. eLife, 3:e01914, 2014.

33. C. A. B. Pearson, S. P. Silal, M. W. Z. Li, J. Dushoff, B. M. Bolker, S. Abbott, C. van Schalkwyk, N. G. Davies, R. C. Barnard, W. J. Edmunds, J. Bingham, G. Meyer-Rath, L. Jamison, A. Glass, N. Wolter, N. Govender, W. S. Stevens, L. Scott, K. Mlisana, H. Moultrie, and J. R. C. Pulliam. Bounding the levels of transmissibility & immune evasion of the Omicron variant in South Africa. medRxiv, pages 2021–12, 2021.

34. R. Earnest, R. Uddin, N. Matluk, N. Renzette, S. E. Turbett, K. J. Siddle, C. Loreth, G. Adams, C. H. Tomkins-Tinch, M. E. Petrone, J. E. Rothman, M. I. Breban, R. T. Koch, K. Billig, J. R. Fauver, C. B. F. Vogels, K. Bilguvar, B. De Kumar, M. L. Landry, D. R. Peaper, K. Kelly, G. Omerza, H. Grieser, S. Meak, J. Martha, H. H. Dewey, S. Kales, D. Berenzy, K. Carpenter-Azevedo, E. King, R. C. Huard, V. Novistsky, M. Howison, J. Darpolor, A. Manne, R. Kantor, S. C. Smole, C. M. Brown, T. Fink, A. S. Lang, G. R. Gallagher, V. E. Pitzer, P. C. Sabeti, S. Gabriel, B. L. MacInnis, New England Variant Investigation Team, R. Tewhey, M. D. Adams, D. J. Park, J. E. Lemieux, and N. D. Grubaugh. Comparative transmissibility of SARS-CoV-2 variants delta and alpha in New England, USA. Cell Rep. Med., 3(4), 2022.

35. R. M. Anderson and R. M. May. Infectious diseases of humans: dynamics and control. Oxford university press, 1991.

36. S. Ballesteros, E. Vergu, and B. Cazelles. Influenza A gradual and epochal evolution: insights from simple models. PLoS One, 4(10):e7426, 2009.

37. E. Volz, S. Mishra, M. Chand, J. C. Barrett, R. Johnson, L. Geidelberg, W. R. Hinsley, D. J. Laydon, G. Dabrera, Á. O’Toole, R. Amato, M. Ragonnet-Cronin, I. Harrison, B. Jackson, C. V. Ariani, O. Boyd, N. J. Loman, J. T. McCrone, S. Gonçalves, D. Jorgensen, R. Myers, V. Hill, D. K. Jackson, K. Gaythorpe, N. Groves, J. Sillitoe, D. P. Kwiatkowski, S. Flaxman, O. Ratmann, S. Bhatt, S. Hopkins, A. Gandy, A. Rambaut, and N. M. Ferguson. Assessing transmissibility of SARS-CoV-2 lineage B. 1.1. 7 in England. Nature, 593(7858):266–269, 2021.

38. H. C. Kung, K. F. Jen, W. C. Yuan, S. F. Tien, and C. M. Chu. Influenza in China in 1977: recurrence of influenzavirus A subtype H1N1. Bull. World Health Organ., 56(6):913, 1978.

39. D. Hannant, J. A. Mumford, and D. M. Jessett. Duration of circulating antibody and immunity following infection with equine influenza virus. Vet. Rec., 122(6):125–128, 1988.

40. D. J. D. Earn, J. Dushoff, and S. A. Levin. Ecology and evolution of the flu. Trends Ecol. Evol., 17(7):334–340, 2002.

41. O. Restif and B. T. Grenfell. Vaccination and the dynamics of immune evasion. J. R. Soc. Interface, 4(12):143–153, 2007.

42. G. Barabás, M. J. Michalska-Smith, and S. Allesina. The effect of intra-and interspecific competition on coexistence in multispecies communities. The American Naturalist, 188(1):E1–E12, 2016.

43. J. M. Levine, J. Bascompte, P. B. Adler, and S. Allesina. Beyond pairwise mechanisms of species coexistence in complex communities. Nature, 546(7656):56–64, 2017.

44. A. Nisalak, T. P. Endy, S. Nimmannitya, S. Kalayanarooj, U. Thisayakorn, R. M. Scott, D. S. Burke, C. H. Hoke, B. L. Innis, and D. W. Vaughn. Serotype-specific dengue virus circulation and dengue disease in Bangkok, Thailand from 1973 to 1999. Am. J. Trop. Med. Hyg., 68(2):191, 2003.

45. B. Bolker and B. T. Grenfell. Space, persistence and dynamics of measles epidemics. Philos. Trans. R. Soc. B, 348(1325):309–320, 1995.

46. V. Andreasen and G. Dwyer. Seasonality and the coexistence of pathogen strains. The American Naturalist, 201(5):639–658, 2023.

47. H. L. Wells, C. M. Bonavita, I. Navarrete-Macias, B. Vilchez, A. L. Rasmussen, and S. J. Anthony. The coronavirus recombination pathway. Cell Host Microbe, 31(6):874–889, 2023.

48. M. J. Mina, C. J. E. Metcalf, A. B. McDermott, D. C. Douek, J. Farrar, and B. T. Grenfell. A Global lmmunological Observatory to meet a time of pandemics. eLife, 9:e58989, 2020.

## References

4. S. Nickbakhsh, C. Mair, L. Matthews, R. Reeve, P. C. D. Johnson, F. Thorburn, B. Von Wissmann, A. Reynolds, J. McMenamin, R. N. Gunson, and P. R. Murcia. Virus–virus interactions impact the population dynamics of influenza and the common cold. Proc. Natl. Acad. Sci. U.S.A., 116(52):27142–27150, 2019.

15. M. Meijers, D. Ruchnewitz, J. Eberhardt, M. Luksza, and M. Lässig. Population immunity predicts evolutionary trajectories of SARS-CoV-2. Cell, 186(23):5151– 5164, 2023.

17. S. Takahashi, Q. Liao, T. P. Van Boeckel, W. Xing, J. Sun, V. Y. Hsiao, C. J. E. Metcalf, Z. Chang, F. Liu, J. Zhang, J. T. Wu, B. J. Cowling, G. M. Leung, J. J. Farrar, H. R. van Doorn, B. T. Grenfell, and H. Yu. Hand, foot, and mouth disease in China: modeling epidemic dynamics of enterovirus serotypes and implications for vaccination. PLOS Med., 13(2):e1001958, 2016.

42. G. Barabás, M. J. Michalska-Smith, and S. Allesina. The effect of intra-and inter-specific competition on coexistence in multispecies communities. The American Naturalist, 188(1):E1–E12, 2016.

49. N. M. Ferguson, C. A. Donnelly, and R. M. Anderson. Transmission dynamics and epidemiology of dengue: insights from age–stratified sero–prevalence surveys. Philosophical Transactions of the Royal Society of London. Series B: Biological Sciences, 354(1384):757–768, 1999.

50. R. Gani and S. Leach. Transmission potential of smallpox in contemporary populations. Nature, 414(6865):748–751, 2001.

51. P. E. M. Fine, Z. Jezek, B. Grab, and H. Dixon. The transmission potential of monkeypox virus in human populations. Int. J. Epidemiol., 17(3):643–650, 1988.

52. S. Wilks. Racmacs: Antigenic Cartography Macros, 2023. R package version 1.2.9, https://github.com/acorg/Racmacs/.

53. F. T. Wen, A. Malani, and S. Cobey. The potential beneficial effects of vaccination on antigenically evolving pathogens. Am. Nat., 199(2):223–237, 2022.

54. R. Sender, Y. Bar-On, S. W. Park, E. Noor, J. Dushoff, and R. Milo. The unmitigated profile of COVID-19 infectiousness. eLife, 11:e79134, 2022.

55. F. Campbell, B. Archer, H. Laurenson-Schafer, Y. Jinnai, F. Konings, N. Batra, B. Pavlin, K. Vandemaele, M. D. Van Kerkhove, T. Jombart, O. Morgan, and O. L. P. de Waroux. Increased transmissibility and global spread of SARS-CoV-2 variants of concern as at June 2021. Eurosurveillance, 26(24):2100509, 2021.

56. K. Sun, S. Tempia, J. Kleynhans, A. von Gottberg, M. L. McMorrow, N. Wolter, J. N. Bhiman, J. Moyes, M. du Plessis, M. Carrim, A. Buys, N. A. Martinson, K. Kahn, S. Tollman, L. Lebina, F. Wafawanaka, J. D. du Toit, F. X. Gómez-Olivé, T. Mkhencele, C. Viboud, and C. Cohen. SARS-CoV-2 transmission, persistence of immunity, and estimates of Omicron’s impact in South African population cohorts. Sci. Transl. Med., 14(659):eabo7081, 2022.

57. COVID-19 Forecasting Team. Past SARS-CoV-2 infection protection against reinfection: a systematic review and meta-analysis. Lancet, 401(10379):833–842, 2023.

58. S. P. Ellner, R. E. Snyder, P. B. Adler, and G. Hooker. An expanded modern coexistence theory for empirical applications. Ecol. Lett., 22(1):3–18, 2019.

